# Near–optimal decoding of transient stimuli from coupled neuronal subpopulations

**DOI:** 10.1101/007450

**Authors:** James Trousdale, Sam Carroll, Fabrizio Gabbiani, Krěsimir Josić

## Abstract

Coupling between sensory neurons impacts their tuning properties and correlations in their responses. How such coupling affects sensory representations and ultimately behavior remains unclear. We investigated the role of neuronal coupling during visual processing using a realistic biophysical model of the vertical system (VS) cell network in the blow fly. These neurons are thought to encode the horizontal rotation axis during rapid free flight manoeuvres. Experimental findings suggest neurons of the vertical system are strongly electrically coupled, and that several downstream neurons driving motor responses to ego-rotation receive inputs primarily from a small subset of VS cells. These downstream neurons must decode information about the axis of rotation from a partial readout of the VS population response. To investigate the role of coupling, we simulated the VS response to a variety of rotating visual scenes and computed optimal Bayesian estimates from the relevant subset of VS cells. Our analysis shows that coupling leads to near–optimal estimates from a subpopulation readout. In contrast, coupling between VS cells has no impact on the quality of encoding in the response of the full population. We conclude that coupling at one level of the fly visual system allows for near–optimal decoding from partial information at the subsequent, pre-motor level. Thus, electrical coupling may provide a general mechanism to achieve near–optimal information transfer from neuronal subpopulations across organisms and modalities.

## 1 Introduction

Flying organisms require fast, reliable feedback regarding ego-motion. This information is extracted from the optic flow – the motion of the external world as perceived by the organism (Lee and Kalmus, 1980; Borst and Bahde, 1988). In the visual system of the fly, neurons of the lobula plate receive as input a two-dimensional, retinotopic representation of the optic flow, allowing them to encode rotational and translational velocities (Borst and Haag, 2002; Borst and Weber, 2011). The lobula plate serves as a primary relay between early vision and downstream motor centers (Strausfeld and Bassemir, 1985; Haag et al., 2007; Wertz et al., 2008).

Approximately sixty large tangential cells responsive to wide-field motion have been identified within the lobula plate of each hemisphere of the blow fly (Hengstenberg, 1982; Hausen, 1984). Ten of these neurons comprise the *vertical system* (VS) which is thought to encode the azimuthal direction of rotations in the horizontal plane (Figure 1A; Krapp and Hengstenberg, 1996). These cells were the focus of our study.

**Figure 1.**
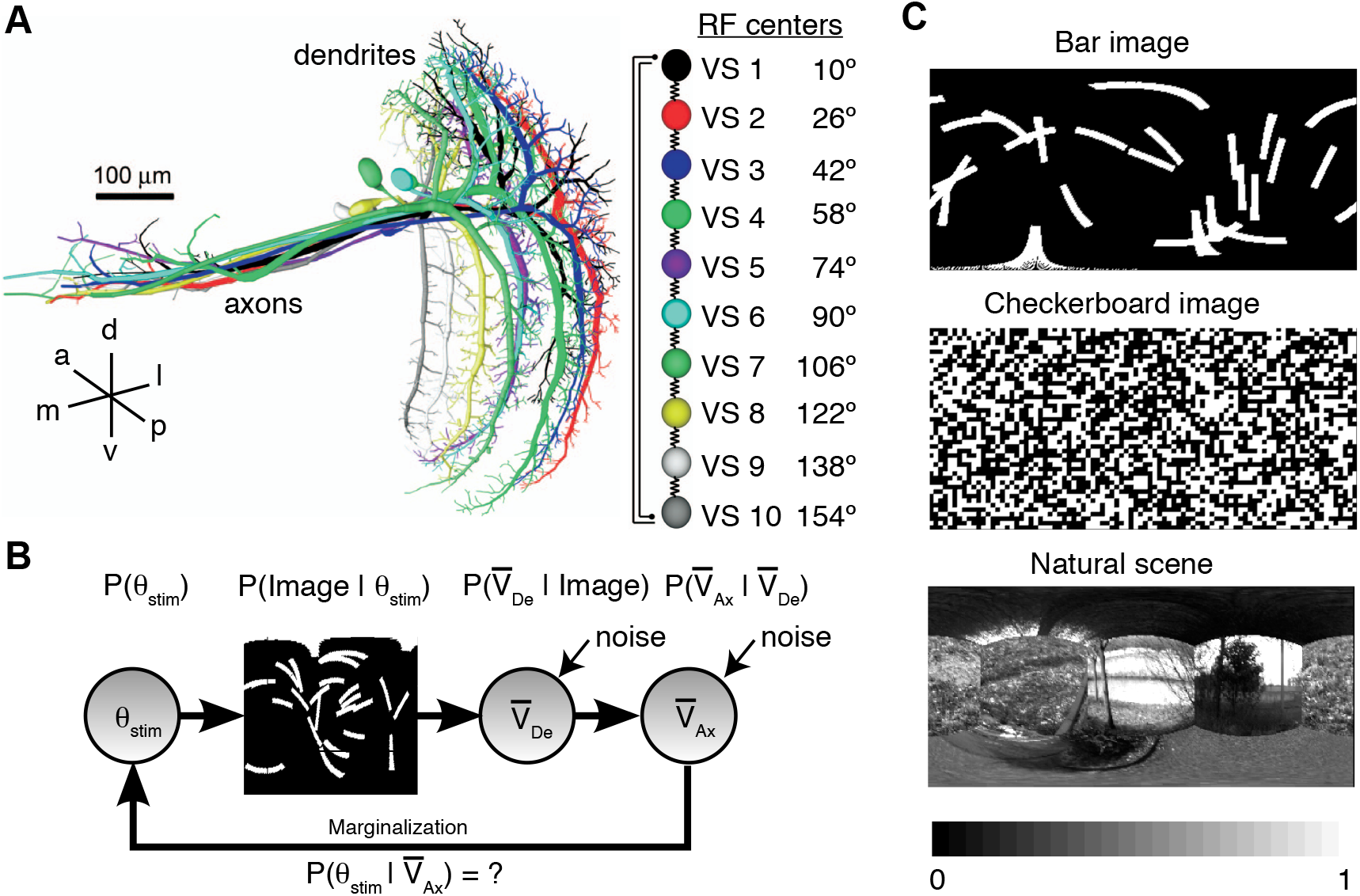
The VS network extracts motion parameters from optic flow-related information. **(A)** (Left) The ten VS cells in one lobula plate (LoP) as reconstructed from two-photon image stacks. Each neuron is T-shaped, with an elongated dendrite sampling a thin vertical stripe in the retinotopically organized LoP (VS 1 to 10 arranged from distal to proximal in the LP, Hengstenberg et al., 1982). Inset indicates approximate orientation (a: anterior, p: posterior, l: lateral, m: medial, d: dorsal, v: ventral). Adapted from Cuntz et al. (2007). (Right) Connectivity scheme of the VS network. VS cell axons are electrically coupled to nearest neighbors. There is a functionally mutually inhibitory (or “repulsive”) interaction between VS1 and VS10. Receptive field (RF) centers indicate azimuthal position in the horizontal equatorial plane of right side VS neurons, taking 0*^◦^* to represent the anteroposterior axis of the fly. Left side VS neuron receptive field centers are given by reflection across 0*^◦^*. **(B)** The marginalization problem: Parameters of ego-motion (such as the axis of rotation, parametrized by *θ_stim_*) are first probabilistically embedded in the external world (Image), and additional layers of variability (noise) are imposed by the processing in VS cells at the dendritic (De) and axonal (Ax) stages (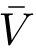 denotes time-averaged membrane potential; see Methods). Reading-out the azimuthal rotation axis from the VS population response amounts to marginalization — extracting the posterior distribution of the stimulus from the axonal responses. **(C)** Example images used to generate the optic flow stimuli presented to the VS network model. Details of image generation are described in the Methods, along with the procedure for generating the rotational optic flow stimuli.

The visual information available to the fly is rich, but only part of it is essential to control flight. The VS cells encode an essential parameter from this complex input – the horizontal axis of ego-rotation. Estimating this parameter is a problem of marginalization, as the quantity of interest must be disentangled from irrelevant visual information (Figure 1B).

Electrical coupling between adjacent VS cells shapes their responses (Haag and Borst, 2004; Farrow et al., 2005). Our goal was to examine the impact of coupling on the representation of the azimuthal angle of rotation in the VS population response. Our results build on previous qualitative observations about the role of coupling (Cuntz et al., 2007; Weber et al., 2008; Elyada et al., 2009).

To examine the role of coupling we presented random rotating images (Figure 1C) as input to a biophysically-plausible model of the VS cell network (Borst and Weber, 2011). In contrast to previous studies, we considered transient responses and applied probabilistic modeling methods to compute optimal Bayesian estimators from VS activity instead of using heuristic or suboptimal estimators.

Anatomical and electrophysiological studies of lobula plate neurons have characterized a pair of pre-motor neurons at the next stage of processing of the fly’s nervous system. The strongest projections of the VS population onto these descending neurons originate from a subset of the VS population (Wertz et al., 2009a). When considering such a partial readout, we found that coupling–induced changes in tuning, correlations and reliability were crucial for an accurate representation of the angle of rotation. Surprisingly, we also found that the quality of the optimal estimate from the collective response of VS cells does not depend on coupling strength.

Gap junction coupling between VS cells can thus impact the accuracy of a subpopulation readout, and hence the fly’s ability to navigate. Our results suggest that across species and modalities electrical coupling can distribute information across a neural population, and significantly increase the performance of estimates extracted from a subpopulation response.

## 2 Materials and Methods

### 2.1 Model of the VS network

Our study is based on a model of the vertical system tangential cells closely related to that of Borst and Weber (2011). We briefly describe the model, and note the differences between the specific implementations. Parameters not explicitly stated, and details of the model not discussed are identical to those given by Borst and Weber (2011).

To mimic stimulation of the fly visual system under a variety of conditions, including natural flight, we started by projecting a random image onto the surface of a sphere. We considered images which consist of randomly arranged bars of varying sizes, as well as random checkerboard images and compositions of natural scenes. Spherical images were rotated about a horizontal axis with varying azimuthal angle, thereby generating a pattern of optic flow (Figure 2A). Details on image and optic flow stimulus generation are given below.

**Figure 2.**
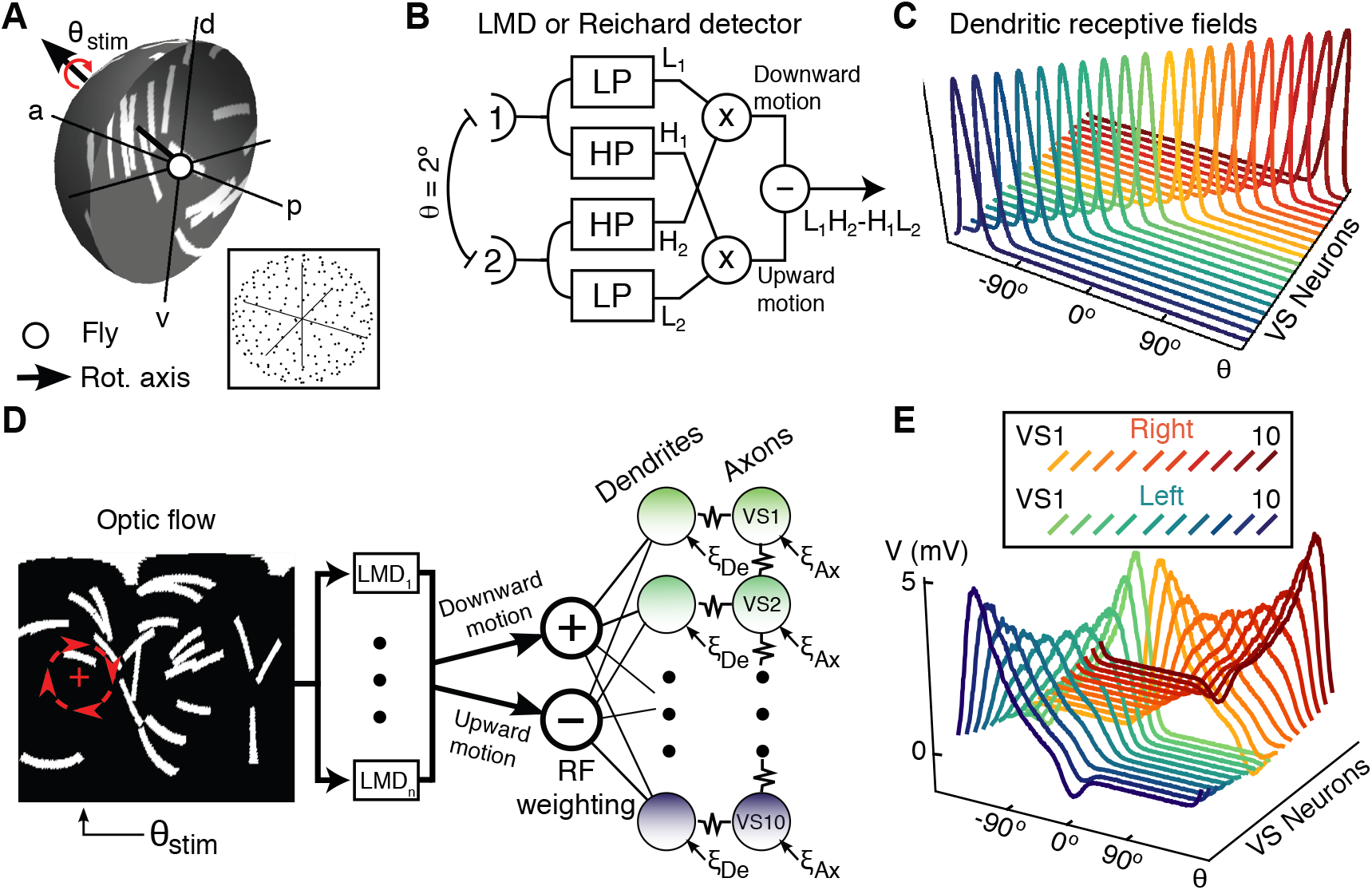
Schematic of the VS network model. **(A)** Spherical image rotation sequences (red curved arrow) were presented to the model fly vertical system (VS). The rotation axis in the equatorial plane is characterized by its azimuth, *θ*_stim_. Lower right inset shows how the Reichardt detectors were arrayed on the surface of the sphere. **(B)** Schematics of the Reichardt detector. LP and HP indicate first order low- and high-pass linear filters, respectively, while *×* and − represent elementary signal multiplication and subtraction steps. Each detector was assembled from two subunits separated by an elevation of 2*^◦^*. The output of the two subunits, maximally sensitive to downward and upward motion, respectively, were fed separately to model VS cells. **(C)** Horizontal cross sections of the dendritic receptive fields for the VS neurons. The frontmost curve is for the left-side VS10 neuron. Proceeding towards the back, blue-green curves correspond first to the left-side VS neurons (decreasing index), then the red-orange curves to the right side (increasing index; see also panel E, upper inset). **(D)** The model is an assembly of a number of components: The optic flow stimulus is generated by rotations of spherical images, and is filtered by the local motion detectors (LMDs). The LMD output is separated into upward and downward components which are mapped to inhibitory (−), and excitatory (+) conductances, respectively, onto the dendrites of the VS neurons. Conductances are weighted by the position of the LMD with respect to the VS cell receptive fields (see C). Resistor symbols indicate electrical coupling of compartments, and *ξ*_Ax_, resp. *ξ*_De_, are independent, intrinsic noise sources to the axons, respectively dendrites, of VS cells. **(E)** Steady-state membrane potential of the twenty coupled VS neurons (*g_gap_* = 1 *µS*) in response to stimulation by a narrow, 10*^◦^* wide horizontal grating with constant downward velocity, centered at angle *θ*. The responses were obtained by sweeping the strip 360*^◦^* around the visual field. Upper inset details color scheme and cell ordering for panels C and E.

Processing of these stimuli by the fly visual system is captured by several successive computational steps. The rotated image sequences (‘optic flow stimuli’) were first filtered by an array of vertically oriented local motion detectors (LMDs or ‘Reichardt detectors’; Reichardt, 1987; Borst et al., 2003; Haag et al., 2004). The LMDs were spaced approximately evenly on the surface of the sphere. There were 5,000 detectors per hemisphere, corresponding approximately to the number of facets on the left and right eyes (Hengstenberg, 1992). The input to a single detector was composed of luminance signals from two vertically-aligned pixels separated by an elevation of 2*^◦^*. First-order filtered low- and high-pass versions of the input from the two pixels were cross-multiplied and then subtracted (Figure 2B). A negative (resp. positive) detector response reflected upward (resp. downward) motion. The downward and upward components were separately weighted according to the dendritic receptive field (RF) of each cell, and activated the dendritic compartment of the VS model neurons as excitatory and inhibitory conductances, respectively. Dendritic receptive fields were vertically-centered Gaussians, with horizontal width 15*^◦^*, and vertical width 60*^◦^* (Figure 2C). Hence each cell effectively sampled the entire vertical surround above and below its receptive field center, given in Figure 1A. Maximal vertical velocity within a cell’s receptive field was generated by azimuthal rotation angles approximately orthogonal to the centers of the receptive field. Such rotations therefore resulted in maximal excitation or inhibition.

The axonal compartments of adjacent, ipsilateral VS neurons are electrically coupled to each other. Figure 2D shows a schematic of the model processing stages. Figure 2E shows the response of each VS neuron to downward stimulation in a narrow (10*^◦^* wide) vertical strip which was swept across the visual field. The vertical strip contained a square-wave horizontal grating with a spatial frequency of 22.5*^◦^* drifting downwards at a constant velocity of 125*^◦^*/ms, corresponding to a temporal frequency of 5.5 Hz.

The axonal and dendritic membrane potentials for the VS neurons in each hemisphere evolve according to:

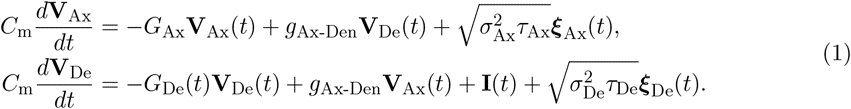

Here **V**_Ax_ and **V**_De_ are vectors whose entries correspond to the 10 axonal and dendritic voltages, respectively. The full VS model consists of two copies of this system, representing the activity of the system in the left and right hemispheres. The two differ only in their receptive field centers (see Figure 1A and Figure 2C). The parameter *g*_Ax-Den_ sets the conductance for the coupling of the axonal and dendritic compartments of each neuron, while *C*_m_ is the membrane capacitance and *g*_L_*_,_*_Ax_,* g*_L_*_,_*_De_ are the leak conductances of each compartment. The membrane time constant of each compartment is *τ_X_* = *C*_m_*/g*_L_*_,X_ , X* = Ax, De. The resting potential of each compartment is zero. Following a perturbation from rest, the membrane potential decays exponentially back to the resting potential with a characteristic timescale *τ_X_* . Intrinsic variability is modeled by standard white noise processes ***ξ***Ax(*t*), ***ξ***De(*t*), and *σ*_Ax_,* σ*_De_ set the noise intensities.

The input currents to the dendrite of cell *i* result in the term *I_i_*(*t*) = *E*_E_*g*_E_*_,i_*(*t*) + *E*_I_*g*_I_*_,i_*(*t*), where *g*_E_*_,i_*(*t*) is the excitatory conductance to cell *i* induced by the optic flow stimulus, *E*_E_ is the associated reversal potential, and likewise for the inhibitory quantities *g*_I_*_,i_*(*t*) and *E*_I_. The matrix *G*_De_(*t*) is diagonal with entries describing the leak conductance, axonal coupling, and input currents,

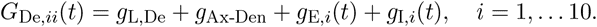

The matrix *G*_Ax_ has entries given by

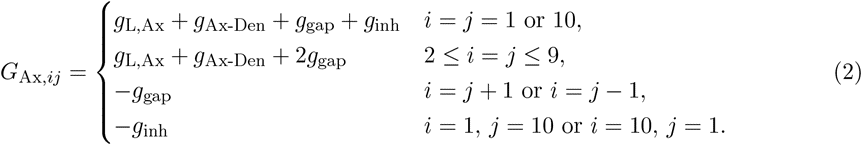

Here, *g*_gap_ sets the strength of the axo-axonal gap junction coupling between adjacent, ipsilateral VS neurons. One difference between our simulation protocol and that of Borst and Weber (2011) is that we generated visual inputs at time steps of 1 ms, but integrated Eq. (1) at a smaller time step of 0.01 ms to guarantee numerical accuracy. We first calculated the conductances elicited by the optic flow stimulus at the coarser time step, then linearly interpolated to obtain a realization of the conductance at the finer timescale. Typical responses of the uncoupled and coupled systems to rotation of a random bar image at *θ*_stim_ = 90*^◦^* are shown in Figure 3A and Figure 3B, respectively. Central to the ability of the VS population to encode the axis of rotation is the strong, non-linear dependence of a VS neuron’s response on the rotational velocity of the visual stimulus within its receptive field.

**Figure 3.**
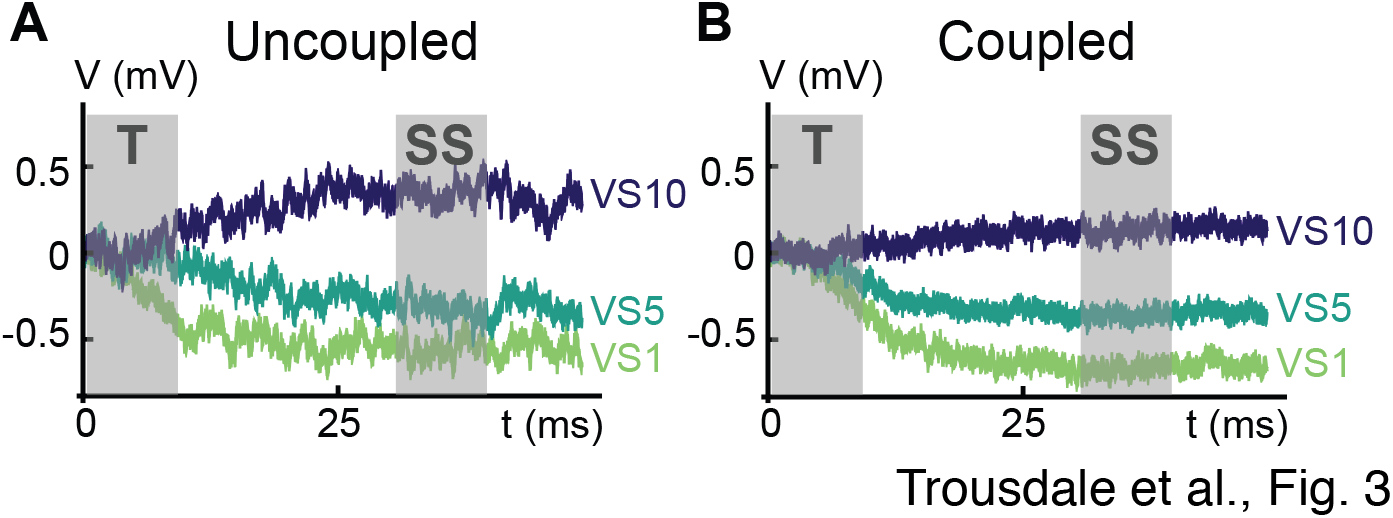
A typical response of the VS network. **(A)** Plot of the temporal response of the left-side VS1, VS5 and VS10 neurons in the uncoupled system (*g*_gap_ = 0 *µS*) to the rotation of a natural scene stimulus (see Figure 1C). Shaded boxes labeled **T** and **SS** indicate time intervals 10 ms in duration over which we average the VS axonal responses to obtain the transient average, 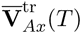 and the steady-state average 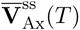, respectively. **(B)** Same as panel A, but for the coupled system (*g*_gap_ = 1 *µS*).

When we consider the encoding of the rotation axis in the VS axonal responses, we take the output of the system to be temporal averages of the axonal membrane potential. For transient responses, the VS output is

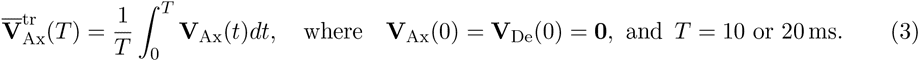

In particular, when considering transient responses, we assume the system starts from rest (0 mV) at the beginning of the period over which we average. Similarly, steady-state responses were computed as

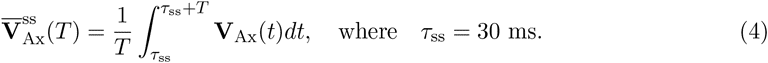

In contrast to the transient response defined above, the steady-state response is defined so that, at the beginning of the integration period (*τ*_ss_), the entire VS system is (approximately) in steady-state. Despite the fast time constants of VS model neurons, they do not immediately reach steady-state, as it takes some time for the motion detector-filtered stimulus received by the VS dendrites to equilibrate. In Figure 3, the shaded boxes indicate the periods over which we calculated the transient and steady-state responses.

It has been observed that there is a mutually inhibitory interaction between the end cells (VS1 and VS10) in each hemisphere. This may be implemented by electrical coupling of VS7-10 to an inhibitory cell Vi which forms a chemical synapse onto the ipsilateral VS1 cell, and electrical coupling of VS1 to an inhibitory cell Vi2 which forms a chemical synapse onto the ipsilateral VS10 cell (Haag and Borst, 2007; Borst and Weber, 2011). Following Weber et al. (2008), we implemented this “repulsive” coupling using a negative-conductance gap-junction between VS1 and VS10 (*g*_inh_ in Eq. (2)). This repulsive coupling was scaled when we changed the strength of axo-axonal gap junction coupling between VS neurons. Unless otherwise specified, we set *g*_inh_ = −0.06*g*_gap_. Our findings do not depend qualitatively on the presence of this connection (results not shown).

For simplicity, we did not model several known functional and anatomical properties of VS cells, such as the rotational structure of their receptive fields (Krapp and Hengstenberg, 1996; Krapp et al., 1998) or dendro-dendritic connections with the dCH neuron (Haag and Borst, 2007). These properties of the VS network are the subject of current experimental investigations (Hopp et al., 2014). Although we do not expect them to significantly affect our conclusions, investigating the impact of such VS cell features is an important avenue for future investigation. Previous computational studies of the VS network have made similar simplifying assumptions (Karmeier et al., 2005; Cuntz et al., 2007; Weber et al., 2008; Elyada et al., 2009, 2013).

### 2.2 Generation **of images and optic flow patterns**

Optic flow patterns were generated by first projecting various types of random images onto the surface of the unit sphere. We considered three classes of random images: random bars, random checkerboards, and natural scenes (see Figure 1C for examples of each type of image). In the first two cases, images were binary — pixel intensities were either 0 or 1 — and for natural scenes, pixel intensities varied continuously between 0 and 1. All images were discretized at 1*^◦^* increments in spherical coordinates. Throughout, images were generated independently across trials.

Random bar images were parameterized by the number of bars, as well as bar width and length. Each bar was generated by first randomly placing an initial line segment of the specified bar width on the surface of the unit sphere. We then expanded this segment along the direction of the length of the bar by rotation about the appropriate axis, turning “on” all pixels along the path touched by the rotating segment. Bar images used in the Results (i.e., Figures 5–6, 7) contained 25 bars of length 40*^◦^* and width 5*^◦^*. Choosing bars with different dimensions or changing the number of bars did not affect the results qualitatively (not shown).

**Figure 4.**
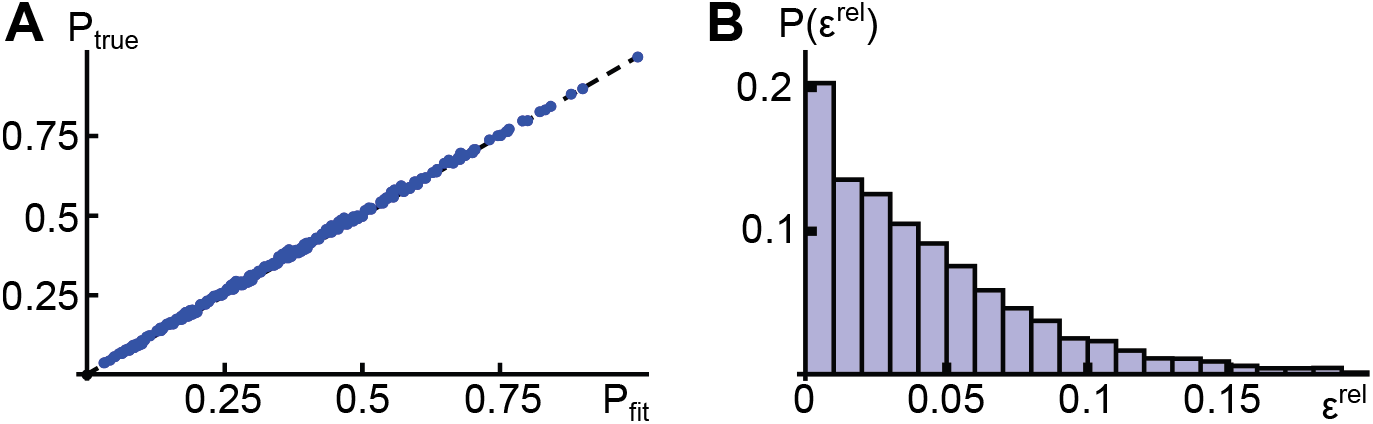
Assessing the copula fit for the transient response distribution. **(A)** Blue points give a P-P plot of the fit copula (*P_fit_*, horizontal axis) against the true, empirical copula (*P_true_*) for a randomly-selected subset of three left-side VS neurons, at a random stimulus angle. We computed the copula probabilities at 1,000 points which divided the unit cube into 1,000 equal sized sub-cubes as described in the text. The black dashed line indicates the diagonal, with agreement between the true and fit models being indicated by the points lying on or near the diagonal. Optic flow presented to the system was generated by the rotation of random bar images, and the copula was fit to the transient response distribution. **(B)** Histogram of relative errors (*∈^rel^*) for copula probabilities. Vertical axis represents fractions of points which lie in the corresponding error range on the horizontal axis. We repeated the simulation of panel A, for a total of twenty random pairings of three left-side VS neurons and stimulus angles. We then computed the relative error (see Eq. (10)) between the true and fit copula probabilities at the 1,000 equally spaced points within the unit cube for all twenty copula fits, and plotted the errors as a histogram.

**Figure 5.**
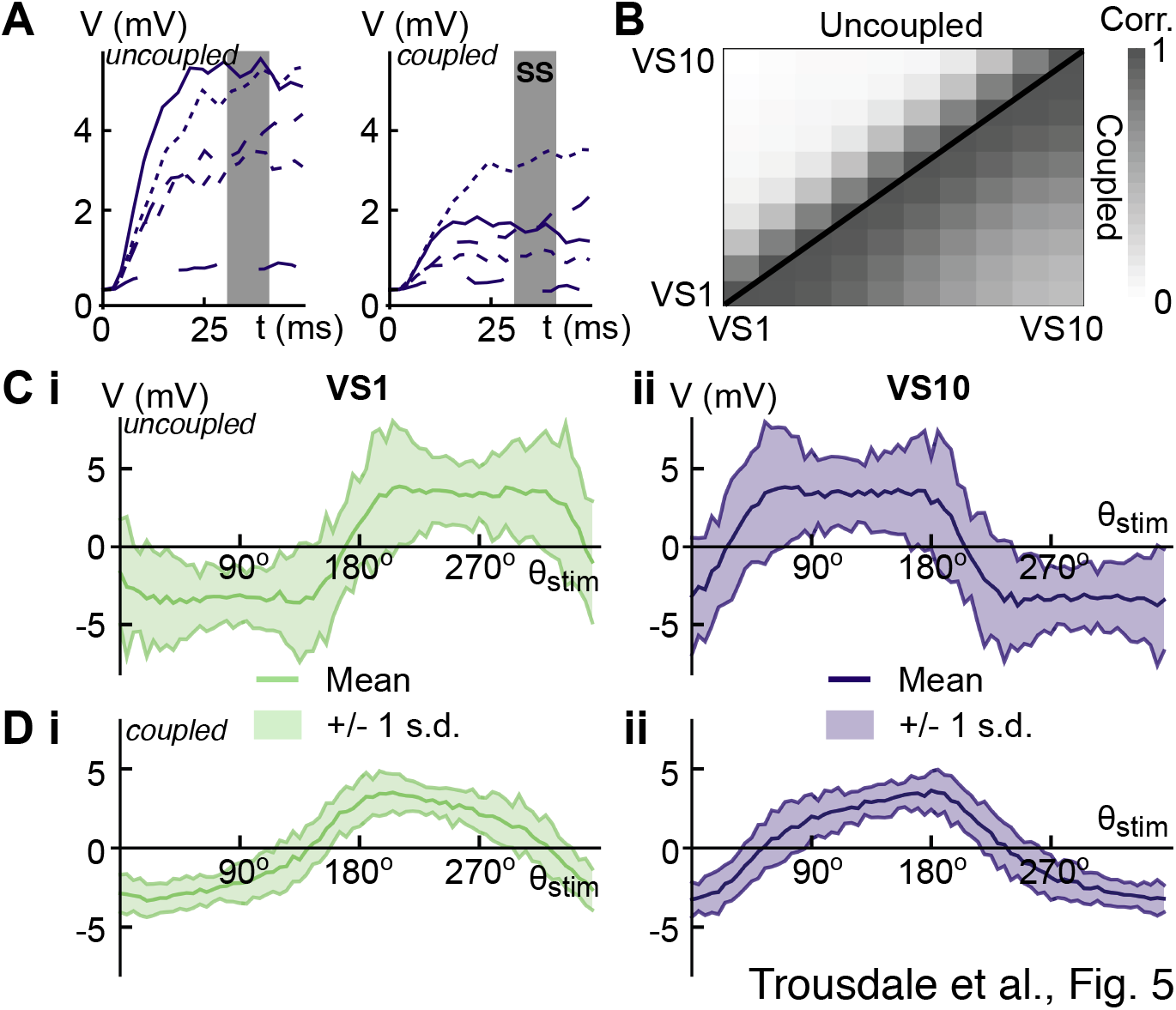
The effect of coupling on VS neuronal dynamics. **(A)** (Left) Typical axonal response of the left-side VS10 cell in the uncoupled network (*g*_gap_ = 0 *µS*) to rotations of bar images about *θ*_stim_ = 90*^◦^*. Different line types indicate different, randomly generated images. (Right) Same as the left panel, but for the coupled system (*g*_gap_ = 1 *µS*). Images rotated to generate optic flow stimuli were the same ones used in the uncoupled system, with matching line types indicating matching image presentations. **(B)** Correlations for the integrated membrane potential in steady-state for the left-side VS neurons. Values above (resp. below) the diagonal are for the uncoupled (resp. coupled) system. Nearby cells were correlated at levels of approximately 0.7 and 0.97 for the uncoupled and coupled systems, respectively. **(C)** Steady-state tuning curve (mean response) and variability as a function of rotation angle for (i) VS1 and (ii) VS10 in the uncoupled system. Shaded areas indicate +/− 1 s.d. of the response distribution. **(D)** Same as C, but for the coupled system. All responses and statistics in this figure were generated in the absence of intrinsic fluctuations (*σ*_Ax_,* σ*_De_ = 0). Stimuli were created by rotations of random bar images (see the Methods).

For checkerboard images, we defined a coarse discretization of the image consisting of 4*^◦^ ×* 4*^◦^* squares, and randomly set all pixels within a square to be zero or one, independently across squares. Lastly, for natural images, we first took a subcollection of one hundred natural scenes from the van Hateren dataset (van Hateren and Schilstra, 1999). These images were selected to exclude man-made objects, sharp edges and gratings. We then randomly selected six (with replacement) of these one hundred images, projected them onto the sides of a cube, which was itself then projected onto a sphere, mimicking the approach of Borst and Weber (2011). For natural scene compositions, we also included an initial rotation of random magnitude about a randomly chosen (not generally horizontal) axis in order to control for the effects of edges between different natural scenes. We also confirmed our results with images chosen randomly from the van Hateren dataset without restrictions. Results were nearly identical (not shown).

The sequences of images comprising optic flow patterns were generated by rotation of the sphere about an axis in the horizontal plane (we did not consider translatory motion). At a given positive time, the value of a pixel was set equal to the value of the pixel obtained by a reversal of the rotation applied to the original image. The rotational velocity was constant across simulations and set to 500*^◦^/s*, falling well within the parameters of typical motion of the fly during flight (Egelhaaf et al., 2012). This value is also consistent with values considered in previous computational studies of the VS network (Karmeier et al., 2005; Cuntz et al., 2007; Weber et al., 2008; Elyada et al., 2009). Increasing or decreasing the rotational velocity to 250*^◦^/s* or 750*^◦^/s* did not affect the results quantitatively, nor change our general conclusions (not shown).

### 2.3 The optimal linear and zero-crossing estimators

To assess the ability of the VS cell network to encode the direction of rotation, we considered several estimators based on the axonal membrane potential of VS cells. In the following three sections, the reader should keep in mind that the random vector v̄ is a surrogate for the time-averaged axonal response of VS cells. We will define the optimal linear estimator of the rotation axis based on the steady-state averaged axonal response of VS cells, 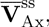 defined in Eq. (4). In subsequent sections, we will also define the minimum mean-square estimator (MMSE) using the transient averaged axonal response, 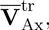 defined in Eq. (3).

For our analysis of the steady-state encoding of the axis of rotation, we applied an optimal linear estimator (OLE), the linear estimator which minimizes the expected value of the squared error over all stimuli and responses (Salinas and Abbott, 1994). We considered a linear, rather than an affine estimator because of the (near) rotational symmetry of the system. The OLE is simple and intuitive, and we used it to demonstrate the impact of gap junction coupling between VS cells. For a more detailed analysis we used the minimum mean-square estimate (MMSE) defined below.

We are interested in estimating the axis of rotation which is characterized by the unit vector, **S**, pointing along its direction. This vector is in the horizontal plane of the fly, with azimuthal angle *θ*_stim_. The optimal linear estimator (OLE) is a linear combination of the responses of the *N* neurons, 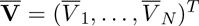 (where *^T^* denotes transposition). To obtain the OLE, we denote the joint probability density of stimuli and responses by 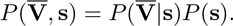 We assume throughout the following that the prior distribution over the stimuli, *P* (**s**), is flat. The tuning curve for the *i^th^* neuron is then

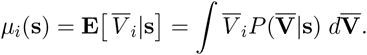

We also set 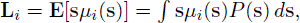, and let *L* = [**L**_1_ . . . **L***_N_* ] be the matrix with columns **L**_1_,* . . . ,* **L***_N_* . We denote the second moment of the responses of the *i^th^* and *j^th^* neurons (averaged across stimulus values) by 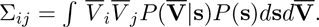 Given an observed response, 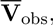 the OLE then has the form

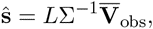

where Σ is the matrix of second moments, Σ*_ij_*. Note that the OLE requires measurement of only first and second moments of the response. It is thus considerably simpler to obtain than the true minimum mean-square estimate (MMSE) defined below which requires knowledge of the full distribution, 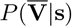. The optimal linear estimator and the MMSE coincide only when the joint distribution 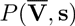 is Gaussian, which is not generally true for the system we consider.

The stimulus **s** estimated by the vector ŝ was a unit vector pointing along the axis of rotation in the horizontal plane. However, the azimuthal angle, *θ*_stim_, of **s** is the behaviorally relevant variable for optomotor control by the fly. We therefore report *the angle θ*̂ *that the vector* ŝ *makes with the direction that the fly is facing*, *i.e.*, the azimuthal angle of ŝ, as in earlier analyses of similar directional stimuli (Lewis and Kristan, 1998; Salinas and Abbott, 1994; Georgeopoulos et al., 1988).

We also implemented an estimator proposed by Cuntz et al. (2007) and Elyada et al. (2009) who suggested that the axis of rotation can be defined as a “zero crossing” of the population response. To obtain this estimator we defined for each cell a *zero angle*. Intuitively, this is the azimuthal rotation angle that is most likely to elicit no response for a given cell. Since rotation about the azimuthal angle coinciding with the center of a VS cell receptive field will elicit little downward and upward motion within the receptive field, the zero angles coincide precisely with the centers of the receptive fields shown in Figure 1A. Then, to estimate the “zero crossing” rotation angle given the response to a rotation about the stimulus angle *θ*_stim_, we search for consecutive pairs of VS neurons which exhibit a sign change in their responses (i.e., their membrane potential responses lie below and above their resting values, respectively). These neurons have the smallest responses to the stimulus, and hence the rotation angle is likely to lie between their respective “zero angles”. The axis of rotation is then estimated by linearly interpolating between the two zero angles based on the responses of these two VS neurons.

More precisely, we define the *zero angle* of the *i^th^* cell on each side, 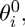 as the angle that maximizes the likelihood of giving a zero axonal response:

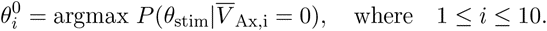

The posterior distribution of the azimuthal angle of the stimulus conditioned on the zero response of a VS cell will generally have two relative maxima, about 180*^◦^* apart. To resolve this ambiguity we choose the angle at which voltage is increasing with increasing *θ*.

To implement the zero-crossing estimator, we first search for a pair of consecutive VS neurons which exhibit responses 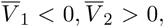 having zero angles 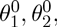 where we assume the two cells are labeled so that 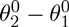 mod 360° ∈ [0, 180∈) A sufficient condition for such a pair to exist is existence of at least one pair of VS neurons exhibiting positive and negative responses, respectively.

Once such a pair is located, the zero-crossing estimate 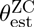 is the angle associated with the zero potential level for the line connecting 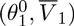 and 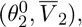

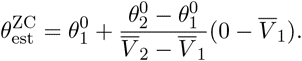

### 2.4 Modeling the joint distribution of VS axonal responses using copulas

For our analysis of the encoding of the axis of rotation in the transient state, we applied the true minimum mean-square estimator (MMSE). Obtaining this estimator of the rotation axis requires estimating the joint probability distribution of axonal membrane potential of VS cells given the stimulus. However, even with the benefit of modern computational power, it is not feasible to directly estimate probability distributions for continuous variates in more than a few dimensions. For this reason, we must first formulate an approximation of the joint probability distribution of VS axonal responses in order to implement the MMSE.

Two approaches to this problem are to fit a maximum entropy distribution that matches a set of empirical statistics of the data (Jaynes, 1957; Roudi et al., 2009; Shlens et al., 2009; Ohiorhenuan et al., 2010; Fairhall et al., 2012), or to apply copulas (Nelsen, 2006). We chose the latter approach, common in the valuation of financial derivatives, but not widely applied in neuroscience (see, however, Berkes et al., 2009; Onken et al., 2009a,b). One advantage of the copula approach is that it allows us to use the empirical marginal probability distributions in the fit.

To fix ideas, we remind the reader that we will fit a copula to the probability distribution of the time-averaged transient VS axonal response, defined in Eq. (3). Thus, we consider a random vector 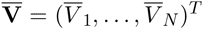 with cumulative distribution function 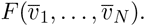 A copula for the distribution function *F* is a function 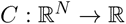 such that

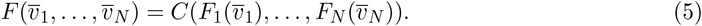

Here *F_i_*(*·*) is the marginal cumulative probability distribution function for the variable 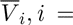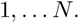. The copula *C* exists for any distribution *F* with marginals 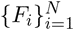 (cf. Nelsen (2006), Theorem 2.10.9). From Eq. (5), it is clear that *C* determines completely the inter-variable dependence structure contained in the distribution *F* in terms of the marginal distributions, *F_i_*.

An *N*-dimensional copula is equivalent to a distribution function on the *N*-dimensional unit hypercube [0, 1]*^N^* with uniform marginals: Define the random vector **U** = (*U*_1_,* . . . , U_N_*) where 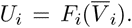 The probability integral transform implies that each *U_i_* is a marginally uniformly distributed random variable (e.g., Gabbiani and Cox (2010), Sect. 11.8). The copula *C*(*u*_1_,* . . . , u_N_*) for 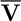 has an equivalent definition as the distribution function of **U**.

The ‘curse of dimensionality’ prevents us from directly approximating the corresponding copula *C*. A common approach is to select a parametrized copula family which can then be fit via the maximum likelihood principle (Yan, 2007). We applied the Gaussian copula (Xue-Kun Song, 2000), which takes the form

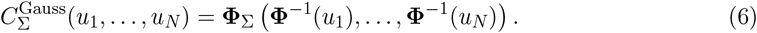

Here **Φ**_Σ_ is the joint Gaussian distribution function with correlation matrix Σ (i.e., Σ*_ii_* = 1 for each *i*) and **Φ** is the standard univariate Gaussian distribution function. Given independent identically distributed samples of a random vector 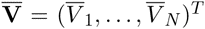 with marginals {*F_i_*}, the correlation Σ*_ij_* equals (Bouýe et al., 2000)

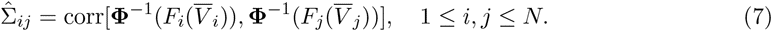

Here corr(*x; y*) denotes the correlation coefficient of *x* and *y* (Xue-Kun Song, 2000).

Given a general copula distribution function *C* as defined in Eq. (5), the copula density is defined by

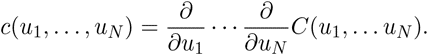

The joint density *f* corresponding to the distribution *F* may be written as

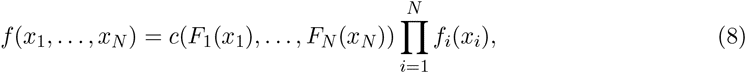

where each *f_i_* is the marginal density corresponding to the distribution *F_i_*. The density of the Gaussian copula may be expressed in closed form as

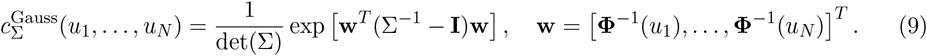

We fit a Gaussian copula to the transient response of the system, 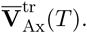. In order to test the goodness of the fit distribution, we selected twenty random subsets of three left-side VS neurons, and associated with each subset a random stimulus rotation angle. We then sampled the marginal copula for each subset (i.e., *C*(*u_i_, u_j_, u_k_*) with 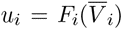 in Eq. (5)), comparing it with the fit copula utilized in the MMSE calculations using Eqs. (6, 7). We compared the empirically observed and fit copula distribution values at 1,000 equally spaced points in the unit cube of the form (0.1*i,* 0.1*j,* 0.1*k*), 1 ≤ *i, j, k* ≤ 10. In Figure 4A, we present a probability-probability (P-P) plot of the true (empirical) copula values against the fit values. In other words, letting *C*^true^ and *C*^fit^ indicate the true and fit copula distribution functions, Figure 4 presents a scatter plot of the 1,000 points

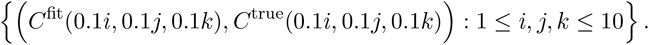

We also computed for each of the twenty subsets and for all 1,000 sample points the relative error between the probabilities from the true and fit copula distributions. In particular, we defined for each point

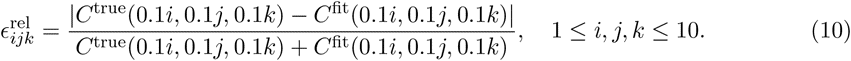

Since the probabilities lie within [0, 1] by definition, the relative errors also lie within [0, 1], with the value 0 indicating a perfect match. In Figure 4B, we plot a histogram of the relative errors for all twenty random subsets (thus comprising a total of 20,000 data points). We found the true and fit copula distributions generally agreed quite well. The average relative error across all 20,000 points was *≈* 0.0438, and 90.3% of relative errors were below 0.1.

### 2.5 Computation of the MMSE

To obtain the MMSE (Kay, 1993), we first simulated the response of the VS network in order to determine an empirical estimate of the marginal distribution functions *F_i_*. We did not assume a parametric form for the marginal distributions, but obtained a discrete estimate by binning values of the membrane potential integral at a sufficiently fine resolution. We then fit the Gaussian copula to the joint responses. Additional details are given at the end of the Methods.

Both the marginal distributions and the copula were determined as a function of the stimulus rotation angle *θ*_stim_ at a resolution of 1*^◦^*. Marginal distribution histograms were approximated from ten thousand samples at each rotation angle, and the copula from one thousand samples. The MMSE was then computed based on 1,600 samples taken at 5*^◦^* increments.

The MMSE of the axis of rotation is calculated as the mean of the posterior distribution of the rotation vector given the axonal membrane potentials. To be precise, given an observed response 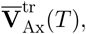 we associated each stimulus angle value *θ*_stim_ with a corresponding two-dimensional unit rotation vector 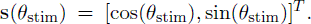 The estimate, 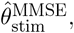 was obtained by first computing an estimate of **s** by averaging over the posterior distribution, i.e.,

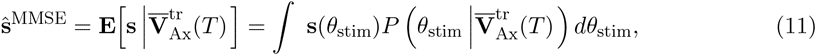

and then reporting the azimuthal angle of this estimate, 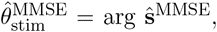 as done above for the OLE and zero-crossing estimators. Values of the posterior density 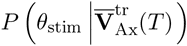 were determined using the fit copula and the measured marginal distributions, along with Eqs. (8, 9). The integral over the posterior density was calculated via simple Riemann integration at a 1*^◦^* discretization.

In much of the Results we obtain the MMSE from a partial readout of the VS response. The estimation procedure is the same regardless of the size of the response vector 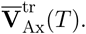 However, we will sometimes use the notation 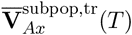 to emphasize that an estimate is based on the response of a subpopulation.

### 2.6 An approximating Ornstein-Uhlenbeck model

Assessing the generality of our results required determining whether our observations depended on the details of the model. In addition, we wished to isolate the essential features governing the modeling results. To do so, we derived a simplified model which shares the essential characteristics of the full model defined in Eq. (1). The first change was to replace the time-dependent parameter *G_De_*(*t*) by a constant, so that the simplified model became an Ornstein-Uhlenbeck (OU) process. Furthermore, we replaced the optic flow generated input by spatially correlated noise. However, cells retained the sinusoidal tuning curves of the full model. Finally, the correlations between the inputs to different cells decayed with the distance between them to capture the effect of overlapping receptive fields. In this simplified model, described by Eq. 12 below, we again changed the magnitude of the diffusive coupling between the cells to examine its impact on encoding.

In form, the corresponding Langevin equations for the evolution of the coupled OU processes **X**_Ax_, **X**_De_ are similar to those of the full model (Eq. (1)):

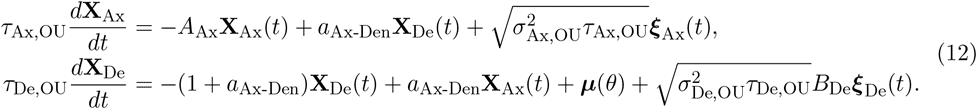

The matrix *A*_Ax_ has entries given by

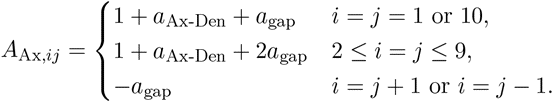

Thus, **X**_Ax_, **X**_De_ are ten-dimensional processes, with a copy of the system for each hemisphere. The two copies are uncoupled, and differ parametrically only in their receptive field centers, as in the full model. We labeled the parameters so as to facilitate comparisons with their counterparts in the full model. For example, the parameter *a*_gap_ captures the coupling between neighboring compartments in the OU model, and corresponds to the parameter *g*_gap_ in the full model.

The tuning curves, ***µ***(*θ*) in Eq. 12, describe the steady-state response in the absence of fluctuations. To approximate the steady-state responses of the full system, we defined individual tuning curves as sinusoids, *µ_i_*(*θ*) = sin(*θ − φ_i_*), where *φ_i_* indicates the receptive field center of the *i^th^* cell.Therefore, a cell will respond most strongly to stimulation angles orthogonal to the cell’s receptive field center, as in the detailed VS model — see Figure 2E. We again obtained estimators from the time integrals of the cell’s responses. In Eq. (12), the last term in the equation for **X**_De_ modeled the correlated input to the dendrites. The matrix *B_De_* was chosen so that correlations to neighboring dendritic compartments decayed exponentially with a space constant of 10*^◦^*: If Δ*φ* was the shorter angular distance between the receptive fields of two neurons, the correlations between the inputs to the two was exp(−Δ*φ/*10*^◦^*).

In Figure 12 we used the following parameters: *τ*_Ax,OU_ = 0.2,* τ*_De,OU_ = 40,* a*_Ax-Den_ = 1,* σ*_Ax,OU_ = *σ*_De,OU_ = 0.2. Receptive field centers, *ϕ_i_,* agreed with the full model. The window of integration for responses was *T* = 10 units of dimensionless time. The coupling strength *a*_gap_ varied, and values used are indicated in the legend of Figure 12.

**Figure 6.**
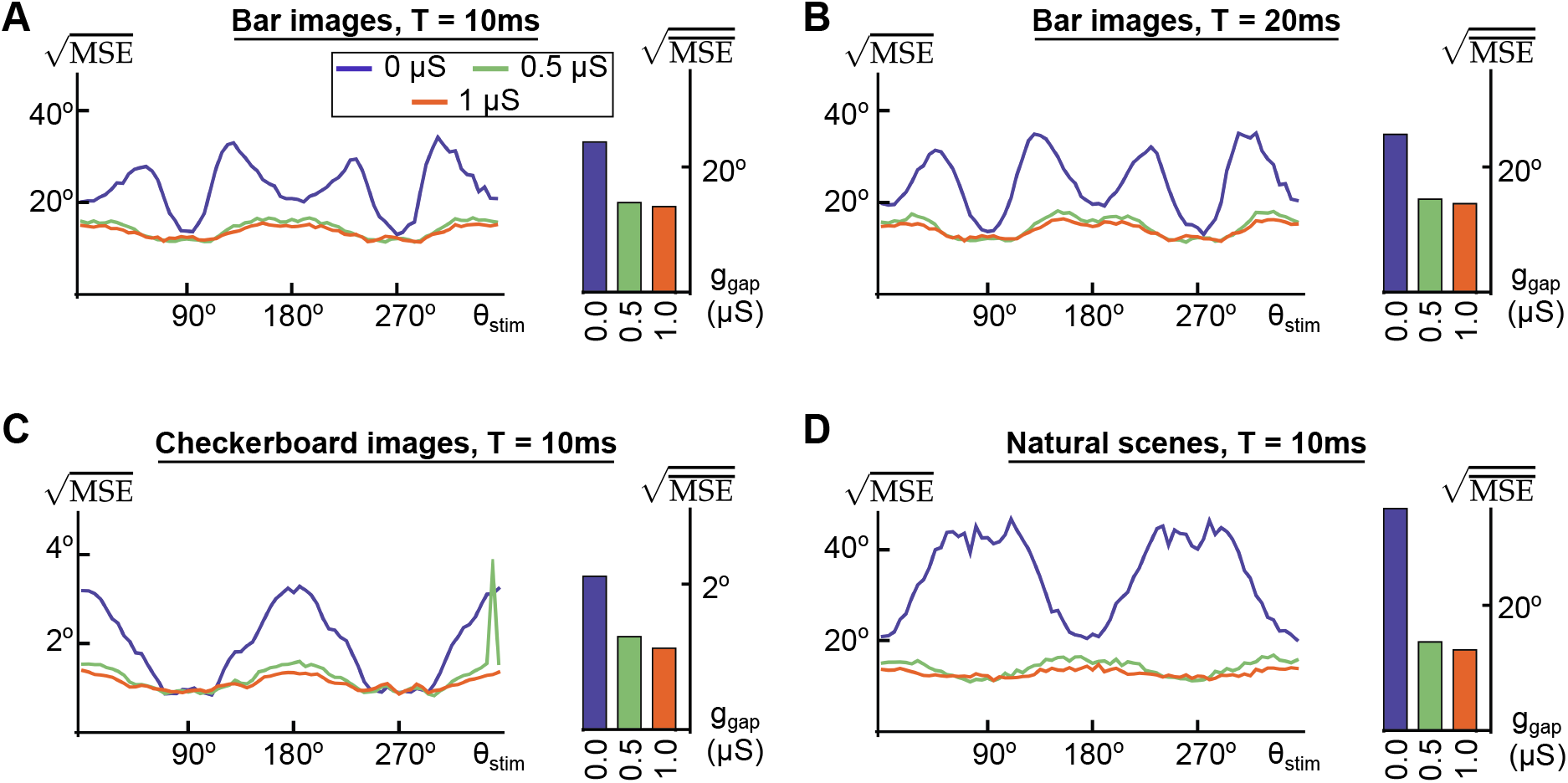
Mean-square error of the MMSE for a partial readout of transient responses. **(A)** (Left) Lines indicate the square root of the mean-square error (MSE) of the MMSE for transient responses to filtered optic flow stimuli generated by the rotation of random bar images with a *T* = 10 ms window of integration. Line colors correspond to different coupling strengths as indicated by the legend. Partial readout refers to the responses of the VS5–7 cells on each side. (Right) Bars represent the square root of the stimulus averaged mean-square error 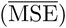 from the data plotted to the left, with bar colors corresponding to line colors. **(B)** Same as A, but for a window of integration of *T* = 20 ms. **(C)** Same as A, but for random checkerboard images. Note the different scaling of the vertical axis. **(D)** Same as A, but for natural scenes.

**Figure 7.**
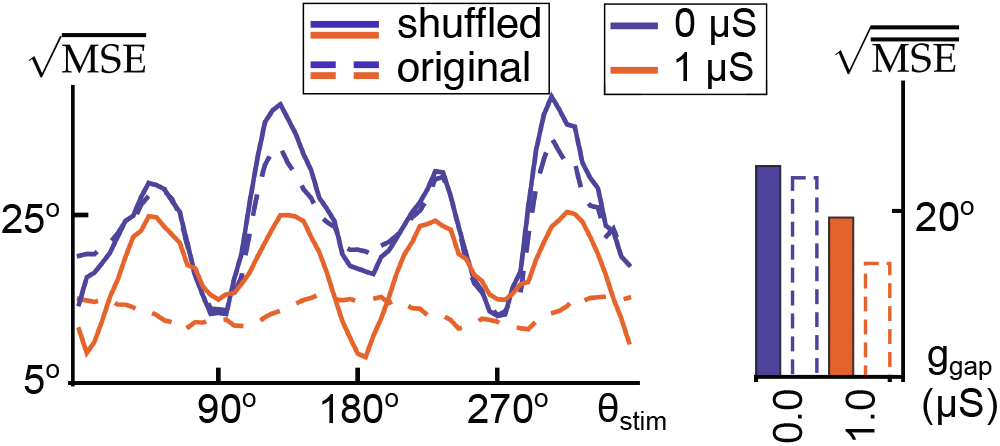
Correlations encode stimulus information in a partial readout. (Left) Solid lines indicate the square root of the mean-square error of the MMSE for trial-shuffled transient responses to filtered optic flow stimuli generated by the rotation of random bar images with a *T* = 10 ms window of integration. Line colors correspond to different coupling strengths indicated (see legend). Dashed lines indicate the error of the partial readout without shuffling for the same strengths of coupling (same data as Figure 6A). The partial readout was formed from the responses of the VS5– 7 cells on each side. (Right) Solid border bars represent the square root of the stimulus averaged mean-square error for the cross-trial shuffled data plotted to the left, with bar colors corresponding to line colors. Dashed border bars indicate the same, but for the non-shuffled data (as in Figure 6).

**Figure 8.**
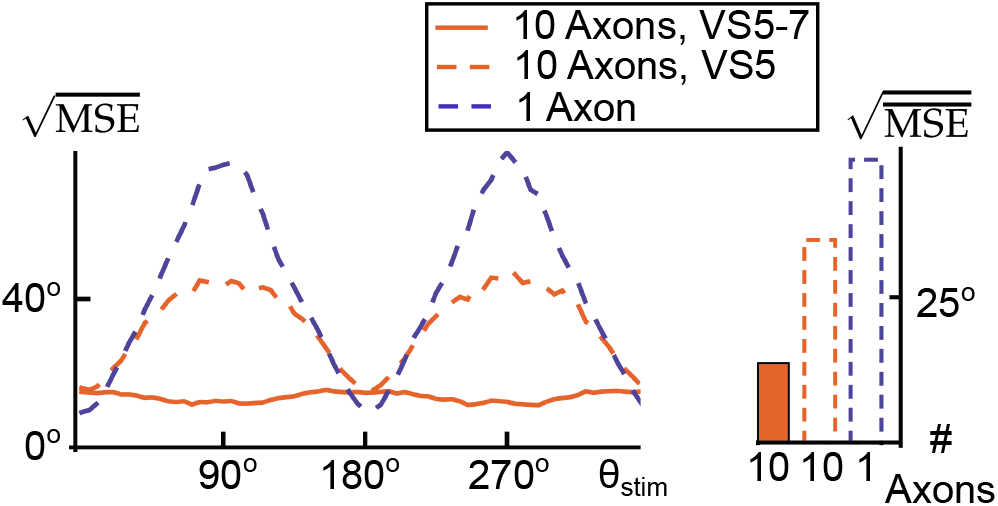
A single cell with a spatially extended dendrite cannot accurately encode the axis of rotation. (Left) Solid orange line indicates the square root of the mean-square error of the MMSE for transient responses to filtered optic flow stimuli generated by the rotation of random bar images with a *T* = 10 ms window of integration, for the full system with a partial readout consisting of the VS5–7 cells in each hemisphere (same data as Figure 6A). The dashed orange line was obtained with the same system as the solid line, with readout from only cell VS5. The dashed blue line indicates the mean-square error for a system consisting of a single axonal compartment in each hemisphere which couples to all ten of the corresponding dendritic compartments, as described in the text. (Right) Bars represent the square root of the stimulus averaged mean-square error for the systems and readouts shown on the left.

**Figure 9.**
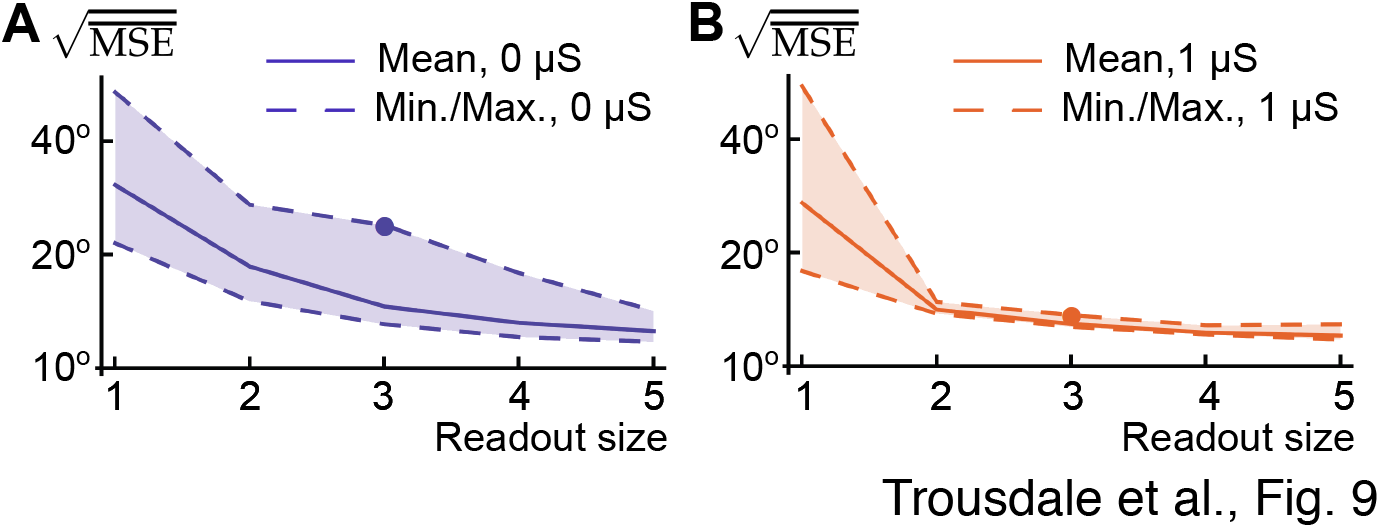
Encoding accuracy depends on cell identities for a partial readout. **(A)** The solid line shows the square root of the average mean-square error of the MMSE calculated from transient responses in the uncoupled system. Input to the system consisted of filtered optic flow stimuli generated by the rotation of random bar images with a *T* = 10 ms window of integration. The mean-square error was averaged across 20 randomly chosen subsets for each readout subset size. For readouts from a single cell, we could only consider ten readouts, corresponding to the ten VS cells. Dashed lines indicate the maximum and minimum values of the MSE observed across the randomly chosen subsets. For size three readouts, we ensured that we included the subset consisting of VS5–7 (the same subset used in Figures 6), and the filled circle indicates the mean-square error for this subset. Note the logarithmic scale of the vertical axis. **(B)** Same as panel A, but for the coupled system.

**Figure 10.**
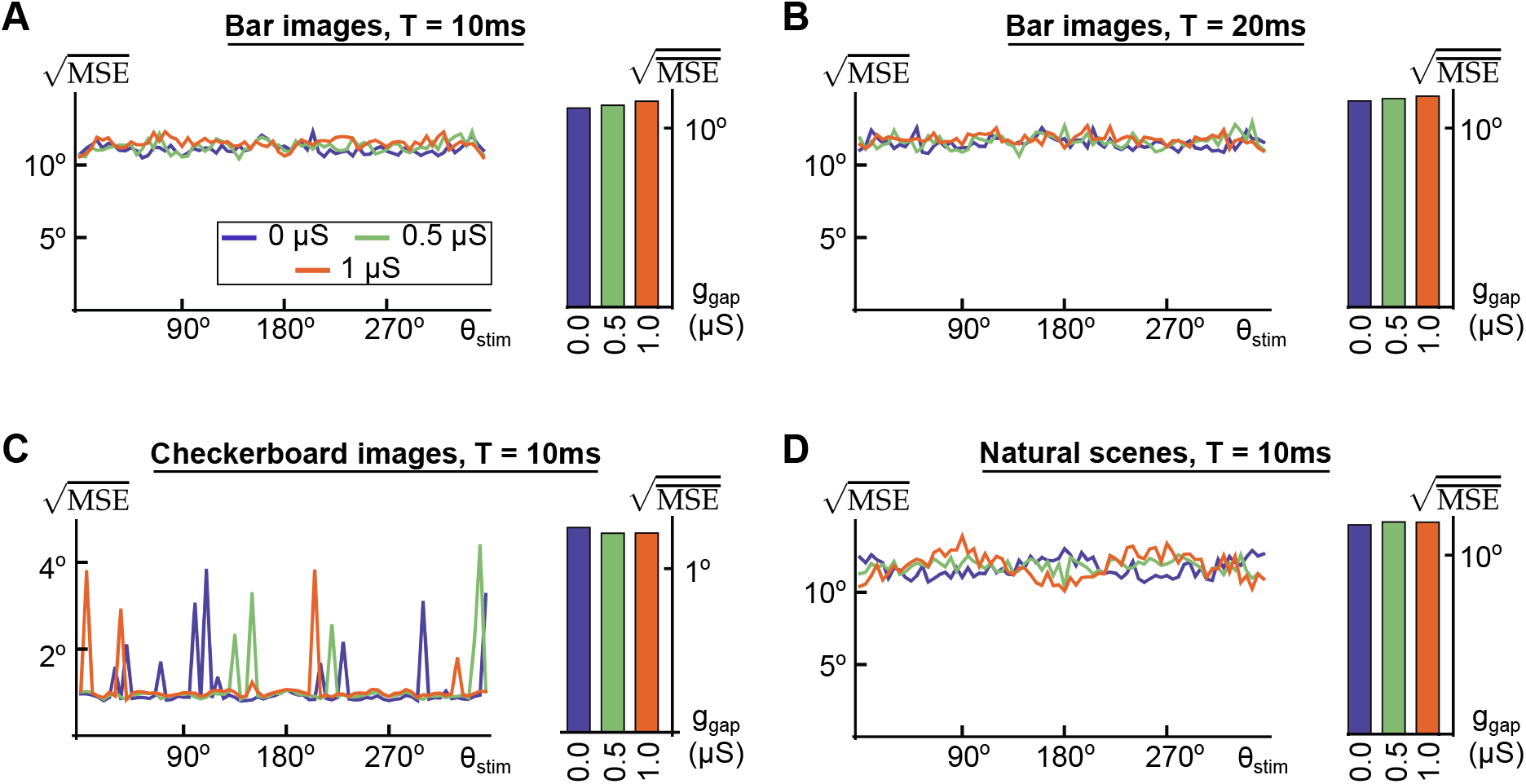
Mean-square error of the MMSE for transient responses. **(A)** (Left) Lines indicate the square root of the mean-square error of the MMSE for transient responses to filtered optic flow stimuli generated by the rotation of random bar images with a *T* = 10 ms window of integration. Line colors correspond to different coupling strengths as indicated by the legend. (Right) Bars represent the square root of the stimulus averaged mean-square error from the data plotted to the left, with bar colors corresponding to line colors. **(B)** Same as A, but for a window of integration of *T* = 20 ms. **(C)** Same as A, but for random checkerboard images. Note the different scaling of the vertical axis. **(D)** Same as A, but for natural scenes.

**Figure 11.**
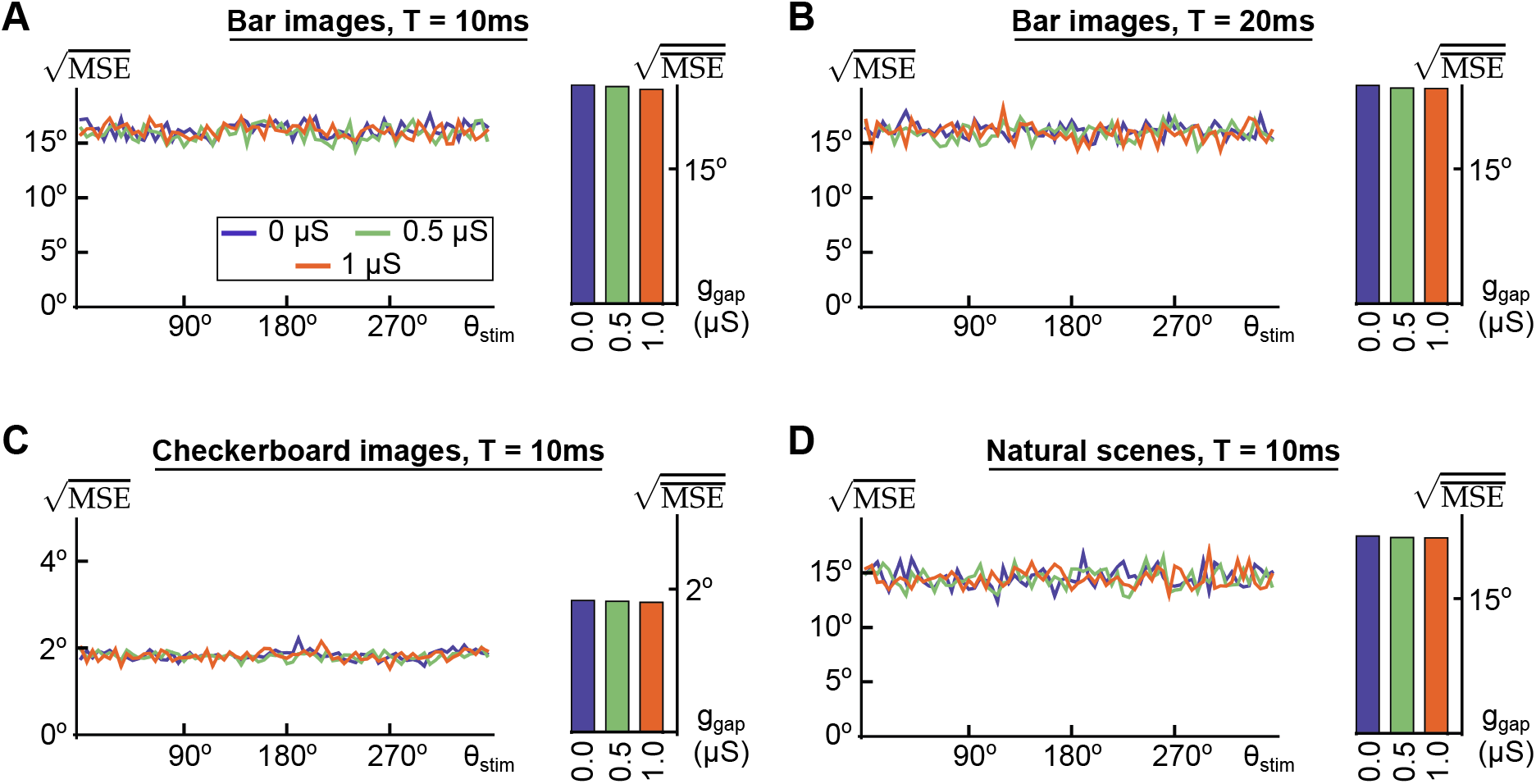
Mean-square error of the OLE for steady-state responses. **(A)** (Left) Lines indicate the square root of the mean-square error of the OLE for steady-state responses to filtered optic flow stimuli generated by the rotation of random bar images with a *T* = 10 ms window of integration. Line colors correspond to different coupling strengths as indicated by the legend. (Right) Bars represent the square root of the stimulus averaged mean-square error for the data plotted to the left, with bar colors corresponding to line colors. **(B)** Same as A, but for a window of integration of *T* = 20 ms. **(C)** Same as A, but for random checkerboard images. Note the different scaling of the vertical axis. **(D)** Same as A, but for natural scenes. Details regarding image generation and the technique for generating the optic flow presented to the model may be found in the Methods.

**Figure 12.**
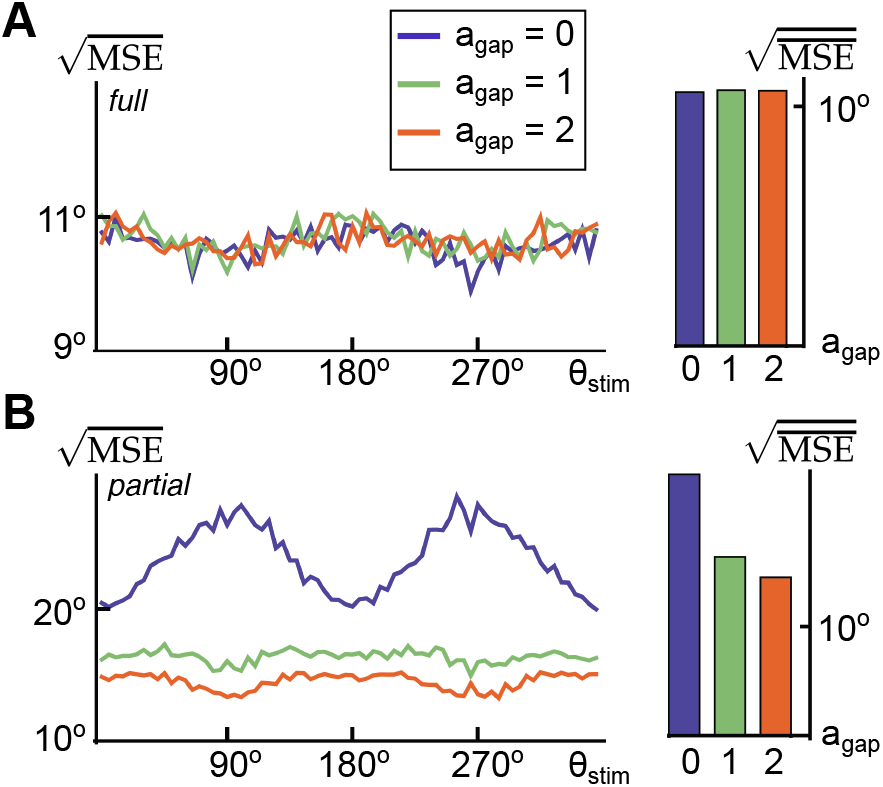
Mean-square error of the MMSE for full and partial readout from an approximating Ornstein-Uhlenbeck system. **(A)** (Left) Lines indicate the square root of the mean-square error of the OLE for transient responses to filtered optic flow stimuli generated by the rotation of random bar images. Line colors correspond to different coupling strengths between “axon” compartments, as indicated by the legend. (Right) Bars represent the square root of the stimulus averaged mean-square error from the data plotted to the left, with bar colors corresponding to line colors. **(B)** Same as A, but for a partial readout formed from the responses of the cells index 5, 6 and 7 on each side, imitating the partial readout considered in the full system (see Figure 6).

### 2.7 Generation of figures

In all figures, to generate a single sample from the true distribution of the VS model axonal response we first used Eq. (1) to model the response of the VS model to random optic flow stimuli. We then computed for each individual simulation the corresponding temporal average (see Eqs. (3) and (4)), to obtain a single sample from the response distribution.

In order to compute the approximate minimum mean-square estimate used to generate Figure 10 and similar figures, we first had to approximate the joint distribution of 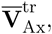 the transient averaged VS axonal responses. We did so in two steps: We first estimated the marginal distribution for each 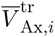 by generating samples as described above and binning at a sufficiently fine resolution. For checkerboard and bar images, we generally binned over the interval [−8 mV, 8 mV] at a resolution of 0.038 mV (420 bins). For natural scenes, where responses were weaker, we generally binned over the interval [−1 mV, 1 mV] at a resolution of 0.0048 mV (420 bins). We verified that bin sizes were small enough so that the results did not change with a further decrease in bin size (results not shown). We determined the marginal histograms at a 1*^◦^* resolution, for each different time window considered. The histograms were approximated using 10,000 data points at each rotation angle. Given this large sample of realizations of the random vector 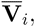 the values 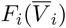 were approximated by applying a rank transformation (Berg, 2009).

In the second step, we fit the Gaussian copula, also by generating samples directly from the model. The correlation matrix which parameterizes the Gaussian copula was determined using Eq. (7). As with the histograms, we fit the copula at each angle at a 1*^◦^* resolution and for each different time window considered. We fit the copula to 1,000 samples generated independently from those used to fit the histograms.

With these approximations of the true histograms and maximum likelihood estimates of the parameters for the fit of the Gaussian copula, we drew 1,600 new samples from the true model, at a spatial resolution of 5*^◦^* for *θ*_stim_. For each sample, we computed the approximate MMSE. By averaging across these samples at each value of *θ*_stim_, we obtained estimates of the relation between *θ*_stim_ and the mean-square error of the MMSE, as plotted on the left of each panel in Figure 10 and similar figures. The averaged mean-square error shown in the bar plots on the right of each panel in Figure 10 was obtained by averaging this data over all values of *θ*_stim_. The averaged mean-square error calculated for each subset in Figure 9 was determined likewise.

Given the intensive nature of the simulations involved in obtaining even a single data point in these figures, all computations had to be performed on supercomputer clusters. For instance, the sum of all computations performed to generate a single curve in the left panel of Figure 10A amounted to over 2,000 CPU hours on 2.2 GHz AMD Barcelona processors. The computer code necessary to implement the VS cell system is deposited in the ModelDB repository accessible at http://senselab.med.yale.edu/modeldb.

## 3 Results

Neurons of the vertical system (VS) in the fly respond to visual input, and encode information about the horizontal axis of ego-rotation (Figure 1). This information is used by cells downstream from the VS system to control flight. We studied how the azimuth of the angle of body rotation is encoded in the response of the VS neuronal network using the model schematically depicted in Figure 2.

Presently, it is not known how downstream neurons read out information from the VS response. However, physiological evidence suggests that only a few VS cells form connections with specific downstream premotor neurons (Haag et al., 2007; Wertz et al., 2008, 2009a,b). We therefore explored the encoding of the axis of ego-rotation in the response of a *subset* of VS cells, by asking what is the best possible estimate of the rotation angle that can be obtained from a partial readout of the VS response. To answer this question, we computed the best estimate (the minimum mean-square estimate or *ideal estimate* from here on) that can be obtained from partial, subpopulation readouts. We found that electrical coupling substantially decreases the error of this estimate. We explain this observation by examining the coupling–induced changes in tuning curves, variability, and correlations of the VS cell population. Surprisingly, we found that coupling has no effect on estimates obtained from a full VS population readout, an observation we go on to explain mathematically. Finally, we validate the generality of these findings using a simplified and abstracted model of the VS network.

### 3.1 Encoding of the azimuthal rotation angle in the VS response

We examined the response of the VS system to a variety of visual stimuli. To simulate the environment surrounding a fly, we first projected images onto the surface of a sphere centered at the fly. The sphere was rotated clockwise about a horizontal axis with azimuthal angle *θ*_stim_, thus generating a pattern of optic flow (Figure 2A). In each trial, we recorded the response of the model VS network to the optic flow generated from a single, random image. Across trials, we used different, randomly generated images: random bars, random checkerboards, and various compositions of natural scenes (see Methods). We show below that, while there were quantitive differences in the VS responses, our results and conclusions hold for all image classes.

Based on our current knowledge of the fly visual system, the optic flow stimulus was filtered by an array of local motion (Reichardt) detectors (Reichardt, 1987; Borst et al., 2003) (Figure 2B). These detectors functionally separated the stimulus into upward and downward motion components which were summed according to the retinotopic receptive field of each neuron (Figure 2C). These signals formed the input to the dendrites of the VS neurons and downward (resp. upward) motion elicited a transient increase in the excitatory (resp. inhibitory) conductance. The magnitude of this conductance change depended non-linearly on the local rotational velocity. Figure 2D provides a schematic of the model.

In the fly, the VS neurons are arranged in a one dimensional array with adjacent, ipsilateral cells coupled via axo-axonal gap junctions (see far right of Figure 2D). The steady-state response of the model network to downward motion in a narrow vertical strip is shown in Figure 2E. Input to one VS cell can impact the axonal response of other cells in the network because of coupling. Thus the ‘effective’ axonal receptive fields of VS cells are much wider than the dendritic receptive fields displayed in Figure 2C. Hence, coupling allows the VS neurons to pool the responses of their neighbors (Cuntz et al., 2007; Elyada et al., 2009). Our goal was to understand the effect of such coupling on the encoding of the azimuth of the axis of ego-rotation in the VS population response.

#### Transient encoding of the axis of rotation in a VS subpopulation

During cruising flight in a stationary environment, flies often move along straight-line segments separated by saccadic periods of rapid rotation. These straight-flight segments occur at rates of up to ten per second and may be as short as 30 ms in duration (Schilstra and van Hateren, 1999; van Hateren and Schilstra, 1999; Boeddeker and Egelhaaf, 2005). Since motor projections of the VS network must pass through intermediate descending neurons (Strausfeld and Bassemir, 1985; Haag et al., 2007; Wertz et al., 2009b, 2008, 2009a), the representation of ego-rotations for compensatory optomotor responses must take place at an even shorter timescale. Similarly, short timescales are likely critical during other natural flying behaviors, such as pursuit and tracking (Land and Collett, 1974; Collett and Land, 1975).

To understand the role of coupling in the VS network we examine its impact on the information available to downstream neurons that are involved in flight control. In particular, we focus on a pair of prominent premotor descending neurons that has been identified within each brain hemisphere. These descending neurons of the ocellar and vertical system (DNOVS) form gap junctions with subsets of the VS cells, and directly innervate motor neurons in the thoracic ganglion of the fly (Haag et al., 2007; Wertz et al., 2008, 2009a,b). DNOVS1 and DNOVS2 couple electrically to ipslateral VS neurons, with the strongest coupling to the VS6-7 and VS5-6 neurons, respectively. The response of these downstream neurons is determined by temporal filtering of the graded response of the VS population rather than instantaneous values of their membrane potentials. We therefore considered time-averaged integrals of the transient response beginning from rest. We denote the vector of these averaged axonal responses by 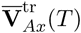 (see Eq. (3) and Figure 3), where *T* denotes the length of the integration window and the “tr” superscript indicates the transient state of the system.

The VS response encodes information about the ego-rotation axis parametrized by its azimuthal angle, *θ*_stim_. We asked what is the best possible estimate of the rotation axis obtainable from the response of neurons that provide input to the DNOVS cells. We therefore computed the estimator that minimizes mean-square error (the ideal estimate) based on the response of the VS5-7 neurons from each hemisphere, and report the corresponding azimuthal angle, 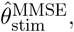 as described by Eq. (11) in the Methods. Computation of 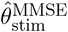 relies on an average taken over the posterior distribution of the stimulus, given the subpopulation VS response, 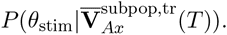 The vector 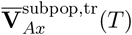 has 6 components corresponding to three cells, VS5-7, in two hemispheres. Later we consider readouts from larger subpopulations, and hence response vectors with more components.

Our model VS network is complex, and it is not obvious how to parametrize the likelihood 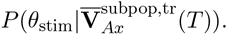 As a result, Eq. (11) cannot be evaluated via Markov Chain Monte Carlo (MCMC) methods, or other techniques designed for efficient sampling from probability distributions (Robert and Casella, 2004). We therefore simulated the model directly to estimate the likelihood, and then computed the integral in Eq. (11) after using copulas to approximate the multivariate distribution of the responses (See Methods for details; Figure 4 illustrates the quality of the copula approximation).

The ideal estimate gives the best possible estimate of the axis of rotation from the responses of a VS subpopulation, 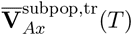. Therefore the mean squared error of any other estimate computed by a downstream population with access to the same VS subpopulation must equal or exceed that of the ideal estimate. While we do not know how the output of the VS system is decoded, our results provide limits on the performance of any postsynaptic decoder.

#### Dynamical effects of coupling on VS network responses

Even in the absence of internal variability, the axis of rotation cannot be decoded perfectly from the VS response. Each visual scene consists of a different arrangement of edges and other features. Thus different scenes rotating about the *same* axis, result in different VS responses. If the parameter of interest, *θ*_stim_, is fixed across a set of trials, we refer to the differences in the VS responses as *overall trial-to-trial fluctuations*. Such fluctuations can be due to variability in visual scenes (external variability), or noise generated internally, but the origin of overall trial-to-trial fluctuations is irrelevant when estimating the axis of rotation from a VS readout.

We illustrate this point in Figure 5 which shows the responses of the VS system to different random bar images. Each image contained the same number of bars of equal shape. However, their arrangement differed from image to image. Even in the absence of intrinsic fluctuations (i.e., *σ*_Ax_ = *σ*_De_ = 0), this resulted in different VS responses when the images were rotated about the same axis (Fig. 5A).

Electrical coupling between VS cells can significantly reduce such overall trial-to-trial fluctuations. Figure 5A shows the effect of coupling using five typical responses of the left-side VS10 neuron to rotations of random bar images about *θ*_stim_ = 90*^◦^* with (left) and without (right) axo-axonal gap junction coupling.

We define the *overall trial-to-trial covariability* in the VS cell responses as the correlation coefficient computed over trials with different visual scenes, but fixed *θ*_stim_. Figure 5B shows the sample (Pearson) correlation coefficient between pairs of VS membrane potentials averaged over the window labeled **SS** in panel A. As expected, correlations increase with coupling. However, even in the absence of coupling, the overlap in the receptive fields of the different VS cells results in strong correlations between neighboring cells.

Figures 5C and D show how steady-state tuning curves (mean responses as a function of *θ*_stim_) and response variability are affected by coupling. In addition to reducing variability, coupling allows the VS neurons to interpolate their responses (Cuntz et al., 2007; Elyada et al., 2009), and increase their orientation coverage (Graf et al., 2011): When coupled to its neighbors, each cell exhibits a graded response to an increased range of stimulus angles. This effect can also be observed by comparing the dendritic receptive fields in Figure 2C with the much broader axonal responses in Figure 2E. Notably, such smoothing takes place without a significant decrease in tuning curve amplitude.

### 3.2 Effect of coupling on VS5-7 subpopulation encoding

What is the impact of coupling on the quality with which the VS5-7 neurons encodes the axis of rotation? The effects described in the previous section point to a potential trade-off: coupling could improve encoding by extending the range of tuning curves, and reducing response variability. On the other hand, increased correlations can decrease the fidelity of a parameter estimate. Whether and to what extent this is the case depends on the specifics of the system (Barlow, 1961; Panzeri et al., 1999; Averbeck et al., 2006b; Shamir and Sompolinsky, 2006; Ecker et al., 2011; Latham and Roudi, 2011).

We examined the impact of coupling on estimating the rotation axis by computing its ideal estimate from a partial readout of the transient VS response to different images and a variable integration time. As shown in Figure 6, coupling greatly increases the accuracy of the estimate obtained from the responses of VS neurons 5–7. Even moderate levels of coupling resulted in a strongly reduced mean squared error of the estimate 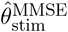 regardless of integration window and type of image presented (bar charts in Figure 6). We will show below that the situation is very different for a full population readout where the mean-square error of the ideal estimate is largely independent of coupling.

### 3.3 Factors determining partial decoding improvement

How does coupling improve decoding? To address this question, we next investigated whether improved estimation of the axis of rotation from the VS5-7 subpopulation is due to changes in individual cell responses. For instance, can changes to tuning curves and the reduction in variability observed in Figure 5C and D explain the improvements in encoding accuracy in Figure 6? Alternatively, can the improvements be explained by changes in the joint response of the VS neurons, such as increases in correlated variability (Figure 5B)?

To obtain a full picture of the factors governing decoding performance, we also investigated how population-level encoding of the axis of rotation depends on the dimensionality of the response: How does a readout from multiple cells receiving distinct dendritic input compare to a readout from a single cell linearly integrating the same dendritic responses? Finally, we asked how the readout from different subpopulations is impacted by coupling.

#### Overall trial-to-trial covariability vs. tuning curve smoothing

To examine the impact of overall trial-to-trial covariability on the error of the optimal subpopulation decoder, we shuffled the responses of each VS neuron across trials, separately for the uncoupled and coupled systems. Specifically, the de-correlated (shuffled) responses were obtained by drawing independently, for each fixed *θ*_stim_, from the set of responses used in Figure 6. This allowed us to maintain the features of the single neuron response to the stimulus (the tuning curve shape), while removing overall trial-to-trial covariability. In the coupled case overall trial-to-trial covariability includes both stimulus-induced and coupling-induced correlations (see above), while in the uncoupled case it consists only in stimulus-induced correlations.

Removing overall trial-to-trial covariability produced a surrogate VS response distribution that differed from the true one. Using this ‘incorrect’ distribution therefore also resulted in a different posterior distribution over the angle *θ*_stim_, resulting in increased ideal estimate error. The magnitude of this increase tells us how important overall trial-to-trial correlations are for the ideal estimate.

As noted above, coupling also smooths the tuning curves and increases coverage. We can compare the ideal estimate obtained in the absence of coupling without shuffling to the ideal estimate obtained with coupling and shuffling. Since overall trial-to-trial covariability is ignored in the second case, any reduction in error is due to changes in the responses of individual cells.

We present the mean-square error of the ideal estimate before and after shuffling in Figure 7. The left plot shows the error as a function of the axis of rotation for two coupling strengths, for trial-shuffled and the original data. The bar chart to the right presents the estimation error for the uncoupled and coupled systems, with and without trial-shuffling.

The response of the VS system is more strongly correlated in the presence of coupling. Therefore shuffling has a stronger effect on the response distribution, and performance of the ideal estimate suffers more when VS cells are coupled. In Figure 7 the difference between the dashed blue bar and the solid orange bar characterizes the impact of tuning curve smoothing due to coupling. The difference in performance between the solid and dashed orange bars characterizes the impact of overall trial-to-trial correlations. Thus, we see that approximately half of the improvement in mean-square error for the subpopulation readout was accounted for by the changes to tuning and the other half by correlated variability: *Changes in the marginal statistics of the response and changes in the correlation structure were of nearly equal importance*.

#### Role of axonal filtering

The VS response is part of a hierarchy of signal processing steps, as shown in Figure 2D. In particular, the axonal system receives input from the dendritic system, which itself has a retinotopic receptive field structure, and also encodes the axis of rotation in its response. We have shown that coupling allows for a more accurate estimate from a subpopulation of the VS cells. However, if only a subset of the VS responses is used by an estimator, why are there ten cells instead of a single cell with a spatially extended dendrite?

To answer this question, we simulated the system with only a single cell (axon) in each hemisphere which received input from all ten dendrites, and left the dendritic structure unchanged. In this case, the axonal response within each hemisphere is univariate and evolves according to (compare with Eq. (1))

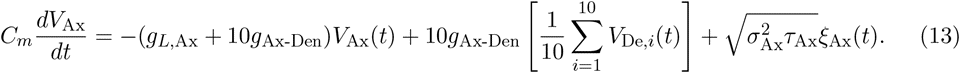

This is equivalent to assuming that a single axonal compartment couples to a single dendritic compartment evolving as the average of the ten dendritic compartments in the full model.

Figure 8 shows that replacing ten cells by one has a strong negative impact on stimulus encoding. Here we compare the error of the ideal estimate for the single axon system to that of the full, coupled system with a partial readout consisting of the VS5–7 cells (as in Figure 6). The average error of the ideal estimate in the full system is smaller by over a factor of four (compare the solid orange and the dashed blue bar, Figure 8, right), and the error is reduced by up to a factor of eight at specific stimulation angles (Figure 8, left).

Thus, multiple cells with differing tuning curves improve the performance of the ideal estimate, even when coupling increases overall trial-to-trial correlations between cells. The improvement is partly due to the additional dimensions available when reading out the VS5–7 responses, compared to a readout from a single compartment. However, this is not all: We compared the performance of the ideal estimate for the single axon system to that of the full, coupled system, with a readout from only the VS5 cell. The coupled system is again superior to the single axon system (compare the dashed orange and the dashed blue line in Figure 8). Hence averaging the responses of all dendritic inputs notably degrades ideal estimate performance. While electrical coupling introduces correlations, responses of the individual VS cells are not identical. A balance between a coupling that allows for the integration of information in a subpopulation, and a distinct response between different VS cells seems best for stimulus encoding in a subpopulation. These results hold for passive dendrites — it is an open question whether a single axon system with active dendrites could perform better.

#### Partial readout subset size and VS cell identity

We asked to what extent our results depend on the particular subset of the VS population used to compute the ideal estimate. Would it matter if the DNOVS cells received input from a different subset of VS cells? To answer this question we randomly selected twenty distinct readouts of sizes up to five (except for readouts of size one, of which there are only ten possible choices). For readouts from three cells, we ensured that the set VS5–7 (the readout considered in Figure 6) was included. For each of the chosen readouts, we computed the mean-square error averaged across all values of *θ*_stim_.

The results are presented in Figure 9. The mean-square error for the coupled network (panel B) decreases rapidly with the size of the readout subpopulation. Improvements are marginal beyond two cells. Strikingly, beyond one cell readouts, there is very little dependence on the particular subset of VS cells on which the estimate is based. In contrast, the average mean-square error across readouts does not decrease as rapidly for the uncoupled network. In this case the mean-square error also depends strongly on the identity of cells in the particular subset. Although the average mean-square error for the uncoupled and coupled network are close for readouts from three or more cells, the error for the “worst” readout is far larger in the uncoupled case.

Despite similarities in membrane and receptive field properties across the VS cells, we observed a large variance in the error depending on cell identity when reading out only a single neuron VS response, for both the uncoupled and coupled network. This variability can be attributed to the unequal coverage of stimulus space by the VS receptive fields, combined with the particular structure of interneuronal coupling. The large variability in the error for readouts from multiple uncoupled cells is due partly to the narrow tuning curves. Uncoupled neurons that are physically close do not communicated with other VS cells. Therefore, the collective ‘orientation coverage’ of uncoupled neighboring cells is lower than for non-adjacent neurons. For such groups of neurons, a larger range of rotation axes will be poorly encoded. In the coupled system, the story is different— when considering a readout from at least two VS neurons from each hemisphere, the identity of the cells is not important. Coupling amongst VS neurons brings about a degree of ‘information democracy’ in which any pair of neurons carries roughly equal information about the axis of rotation.

Overall, coupling was uniformly beneficial to the fidelity of encoding of the axis of rotation in the axonal response. Strikingly, however, the error for the readout of the VS5–7 cells was highest amongst all tested readouts for both the coupled and uncoupled system. Two key factors contributed to this result: first, single cell readouts of each of these neurons yielded a larger error than single cell readouts from other VS neurons (results not shown). In addition, these three cells are physically close, and thus lead to larger estimation errors, as noted above. This phenomenon is closely linked to observations of changes in orientation coverage with population size recently reported in pools of orientation selective cortical neurons used to discriminate sinusoidal gratings drifting in different directions (Graf et al., 2011). Thus, if biological constraints dictate that the rotation angle be estimated primarily from the response of the VS5–7 neurons, coupling of the VS axonal responses becomes of great importance (Figure 6).

Comparing the trends of the mean-square error in Figure 9 in the absence (A) and presence of coupling suggests that coupling has a diminishing effect as subpopulation size increases. We next examine this observation further.

### 3.4 Estimation **of the axis of rotation from the full population response**

We have shown that coupling significantly improves estimates from a partial readout of the VS response. How does coupling affect a readout from the entire VS population?

The performance of the ideal estimate for the angle of rotation estimated from the integrated axonal potentials of the *full* VS population is shown in Figure 10. The error of the ideal estimate depended on image statistics, but surprisingly, it was approximately independent of coupling strength.

As shown in Figure 5, coupling has a strong effect on correlations and tuning curves, and decreases the error of the ideal estimate obtained from a partial readout. It is therefore surprising that the same changes in tuning curves and correlations have no effect on the error of the ideal estimate obtained from the full population response. To understand this difference, we first apply the *optimal linear estimator* to steady-state VS responses (OLE; see Methods). While less general than the ideal estimate, the OLE is easier to analyze. We show that the insights obtained from the OLE in steady-state carry over to the ideal estimate both in steady-state and transient states.

#### The mean-square error of the OLE is independent of coupling in steady-state

We defined the steady-state response of the system of VS cells using averages of the graded responses of the VS population, 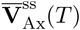 (See Eq. (4)). Here *T* indicates the time window of integration and the superscript “ss” indicates that the system is in steady-state. We then computed the OLE based on 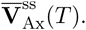

The mean-square error of the OLE as a function of *θ*_stim_ is shown in Figure 11, for three types of images and two integration time windows: Across image types and integration time windows, the mean-square error of the OLE in steady-state was also independent of the strength of axo-axonal coupling. The statistics of the optic flow generated by images from a given class set the baseline level for the mean-square error. However, the error was independent of coupling strength for all classes of images tested. We also verified this for different parameters of the random bar images (results not shown — see the Methods for details of image generation).

#### Explanation of coupling-independence

We first note that since the axonal and dendritic compartments were coupled electrically, the dendritic response was affected by the strength of the axo-axonal coupling. However, the impact of coupling on the vector **V**_De_(*t*) of dendritic responses was limited to a multiplication of the response by a diagonal matrix (result not shown). Except for this scaling, the time course of the dendritic response is dominated by the synaptic input arriving through the visual pathway which reflects the response to the filtered optic flow stimulus. Therefore, to a good approximation, the system may be viewed as hierarchical (Elyada et al., 2009): The motion detector-filtered optic flow stimulus drives the dendrites, and the dendrites drive the axonal compartments, with the activity at each step determined completely by the response at the preceding stage (along with any intrinsic noise sources).

As we show next, a consequence of this observation is that the vector of average axonal responses 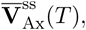 is linearly related to the vector of average dendritic responses 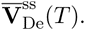 The specific relation between the two vectors changes with coupling. However, because the relation is linear, the axonal and the dendritic response will give the same estimate of the axis of rotation, regardless of coupling. We next make these intuitive observations more precise.

Disregarding intrinsic noise in the system, we have

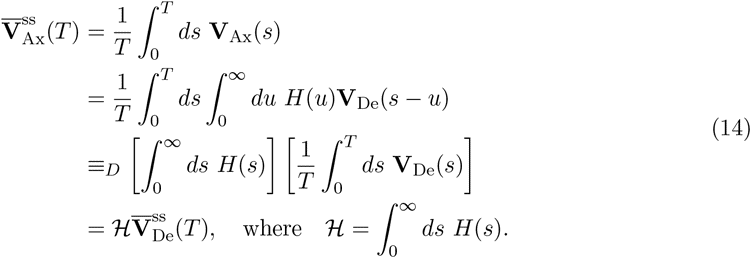

Here, *≡_D_* indicates equality in distribution, and 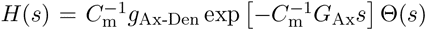 is the exponential filter the axonal system applies to the dendritic response. We note that at positive times, when the Heaviside function Θ(*s*) = 1, *H*(*s*) is a matrix exponential. That the axonal response can be represented by a convolution of a matrix exponential with the dendritic response (second equality) is a general mathematical property of linear systems of differential equations such as those which describe the evolution of **V**_Ax_ (see Eq. (1)).

The equality in distribution in Eq. (14) arises from switching the order of integration and using the time-shift invariance of the dendritic membrane potential distribution under the steady-state assumption. In this case the distributions of **V**_De_(*s − u*) and **V**_De_(*s*) agree for all finite *u*, and the two quantities can be exchanged under an equality in distribution. The dendritic average 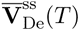 is defined analogously to the axonal quantity in Eq. (4).

Eq. (14) shows that, under the hierarchical assumption, the axonal activity is conditionally independent of the input given the dendritic activity: If the linear relation between axonal and dendritic responses is invertible, the posterior distribution of the stimulus given the axonal response agrees exactly with the distribution conditioned on the dendritic response. In other words, since 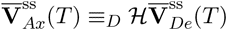 for some invertible matrix *ℋ*, it follows that

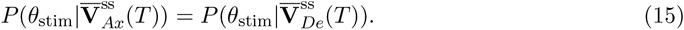

In this situation, there is no change in information about the rotation angle due to coupling. This equality holds as long as 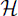 is invertible. Realistic gap junction coupling strengths change the entries in the matrix 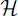, but do not impact its invertibility.

We found that the dendritic responses were independent of coupling up to an invertible linear scaling factor, implying that the posterior distribution 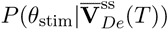 is likewise independent of coupling. Thus, not only will the performance of the OLE be unaffected by coupling in this case, but the same conclusion holds for more general probabilistic estimators (including the ideal estimate considered below, all Bayesian estimators and the maximum likelihood estimator).

#### Independence of the ideal estimate on coupling in the transient state

The explanation of why the ideal estimate error is independent of coupling in the transient state is largely identical to that provided in the steady-state case (see Eq. (14)), with one crucial difference: The equality in distribution on the third line of Eq. (14) does not hold exactly for transient responses. *A priori* it is not obvious that the equality should hold even approximately. However, if the axonal responses are fast then the equality will hold to a good approximation: Intuitively, a fast response means that the axonal filter, *H*(*u*) is not negligible only for *u* close to 0. Since the dendritic voltage changes are relatively slow, **V**_De_(*s − u*) *≈* **V**_De_(*s*) is nearly independent of *u*.

More precisely, the characteristic response timescales of the VS axonal compartments (and of the system filter *H*) are given as the product of the membrane capacitances with the inverses of the eigenvalues of *G*_Ax_. If these eigenvalues are large enough, the equality in distribution in Eq. 14 does hold nearly exactly owing to a separation of the timescale of the axonal response from that of the output integration window. In this case, there again exists an invertible linear relationship between the transient axonal average 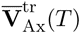 and the corresponding dendritic average 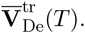.

The VS axonal compartments have baseline time constants on the order of a millisecond (Borst and Weber, 2011). The effective time constants will thus be even smaller, owing to the axo-dendritic and axo-axonal gap junction synapses (Rudolph and Destexhe, 2003). These short integration time constants allow the system to reliably implement a linear transfer from the dendritic to the axonal averages for transient responses, as in Eq. (14). To verify this, we carried out linear regression analysis of the dependence between the axonal and dendritic responses. We found *R*^2^ values for individual coordinates were above 0.999 for all coupling values (*g*_gap_ = 0, 0.5, 1 *µS*) and integration time windows tested (*T* = 10, 20 ms) when the intensity of intrinsic noise was set to zero (results not shown). This indicates that transfer is nearly linear in each axonal dimension. The linear regression was performed for random values of *θ*_stim_, indicating the independence of this transfer matrix from the stimulus value.

In short, the axonal network uses a very fast system filter to institute a highly reliable linear transfer of the averaged transient dendritic response to the average transient axonal response. As in the steady-state case, the entries of the transfer matrix depend on coupling, but its existence and invertibility do not. Hence, the posterior distribution of the stimulus conditioned on the axonal response is nearly identical to that conditioned on the dendritic response (see Eq. (15)), and the latter is approximately independent of coupling (Elyada et al., 2009). As a result, the estimates (and error) of any probabilistic estimators will not depend significantly on the strength of the axo-axonal gap junction coupling within the VS system.

#### Coupling improves partial decoding in a simplified model

We next asked whether our conclusions depended on the details of the model fly visual system we used in our simulations, the statistics of the images presented, or other choices we made in our analysis. For this purpose, we examined whether similar conclusions can be obtained in a simplified, analytically tractable approximation of the full VS model (see the Methods). In this simple Ornstein-Uhlenbeck (OU) system we discarded the Reichardt detectors, and the complex visual input used in the full model. Instead, the input to the system was white noise which was correlated in space to model VS cell’s receptive field overlap.

The ten dendritic and ten axonal compartments were modeled by linear differential equations which shared two essential characteristics with the full model described above. First, we assigned the dendritic compartments tuning curves (i.e., mean responses in the absence of fluctuations) which were sinusoidal functions of the stimulus angle in order to emulate the retinotopic response properties of the VS neurons. Second, the axonal compartments were diffusively coupled.

In this model, we emulated the relatively slow timescale of the dendritic input arriving to the axon terminals, by assigning the dendritic compartments a time constant which was an order of magnitude larger than that of the axonal compartments. Details of the model are given in the Methods.

In this setting, we qualitatively replicated our earlier findings: The performance of the ideal estimate obtained from a partial readout of the axonal responses increased with coupling. With a readout from the entire population, performance was independent of coupling (Figure 12). We did not tune the model, and the result held over a wide range for all parameters (results not shown).

## 4 Discussion

Organisms need to rapidly extract information about ego-motion from complex patterns of optic flow (Figure 1C). We used a simplified, but biophysically realistic model of the fly vertical system (VS) to examine the role of electrical coupling in encoding the azimuthal axis of ego-rotation. We have shown that this parameter can be quickly and accurately extracted from the transient response of a VS subnetwork.

The impact of coupling and correlations on encoding in neuronal populations has been studied extensively (for a review, see Averbeck et al., 2006a). Interestingly, we found that coupling did not affect the error of optimal estimates from complete population responses. The posterior distribution over the azimuthal angle of rotation was unaffected by coupling. Hence, Bayesian or maximum-likelihood estimators were similarly unaffected.

Physiological evidence suggests that part of the VS network drives the response of two of its postsynaptic neurons. The DNOVS 1 and 2 are efferent targets of VS cells projecting to thoracic ganglia where they contact neck motor neurons involved in head stabilization during flight (Haag et al., 2007; Wertz et al., 2008, 2009a). Each DNOVS neuron is most strongly coupled to two VS cells, and the DNOVS pair is coupled predominantly to a subset of three VS neurons in each hemisphere.

Coupling between VS cells significantly improved encoding accuracy of the rotation angle in the response of this VS subpopulation. For this partial readout, the transfer from dendritic to axonal membrane potential is approximately linear, but not invertible. Hence, the performance of the estimator depends on coupling. The dynamical changes induced by coupling resulted in a sub-population readout that could be as accurate as a full population readout. Coupling had the greatest impact on the readout from VS5–7 and post-dendritic processing enabled by the coupling of distinct axonal compartments was crucial to encoding accuracy. These results are robust and general: Model details such as image features and integration timescales had only a quantitative impact.

Based on these results, we can make concrete predictions that can be tested experimentally. These predictions rely on the ability to ablate a single VS cell that does not provide direct input to DNOVS neurons, or block its gap junctions. Under such conditions, the decoding of rotation axes from postsynaptic DNOVS neurons should be differentially affected, depending on which VS cells the rotations preferentially stimulate. If such specific silencing of a VS neuron were possible in the intact animal, we predict a similar differential effect on behaviors controlled by DNOVS neurons.

The fly visual system is an established model for the study of optimal motion encoding and decoding (Laughlin, 1981; Bialek et al., 1991; van Hateren, 1992; Gabbiani and Koch, 1996; Van Steveninck and Laughlin, 1996; Fairhall et al., 2001). To the best of our knowledge, our work is the first to examine the impact of electrical coupling on optimal population coding by using a mathematical and computational analysis of the response of a detailed model of the VS network (Borst and Weber, 2011). Previous arguments about the benefit of coupling were primarily heuristic (Elyada et al., 2009), and generally concerned steady-state responses (Cuntz et al., 2007; Weber et al., 2008). By considering dominant eigenmodes, Weber et al. (2008) showed that coupling leads to a reliable, lower dimensional representation of VS activity. Rotation about a given azimuthal axis results in depolarization of the VS cells located to one side of the axis, and hyperpolarization in the cells located on the opposite side. This prompted Cuntz et al. (2007) and Elyada et al. (2009) to propose estimating the axis of rotation by interpolating the responses of the VS cells, and finding a zero-crossing in the mapping between these responses and the VS cells’ preferred axes of rotation. Coupling reduces response variability, and hence the error of this estimate.

When we implemented such a zero-crossing estimator (see Methods) and compared it to the ideal estimate, we found it to be suboptimal. The estimator usually struggled to achieve reasonable encoding accuracy when presented with natural scenes. Responses to natural scenes were weaker than those induced by the other image types we considered, and the zero-crossing estimator is very susceptible to noise. Furthermore, coupling could either improve or degrade the estimates, depending on image statistics. The zero crossing estimator from a subpopulation response performed even worse, producing errors several times that of the ideal estimate.

Furthermore, it is unclear how the zero-crossing and other proposed suboptimal estimators could be implemented downstream from the VS cells. In contrast, the ideal estimate establishes a baseline for the performance of any estimator. While suboptimal estimators may be affected by coupling in different ways, we found consistent results for the ideal estimate across stimulus conditions. Although neuronal networks may not process information optimally (Loeb, 2012), evidence for approximate optimality exists both at the behavioral and neural level (e.g., Ernst and Banks, 2002; Fetsch et al., 2012). We used optimal estimators as an operational benchmark, revealing the potential capabilities of the system under realistic assumptions.

Two studies are conceptually close to our work: Karmeier et al. (2005) took a Bayesian approach to quantify the encoding efficiency of the axis of rotation in the VS population response. They also proposed time integrals of the VS membrane potentials as readout variables, and examined the impact of population size on encoding in the VS population. We note several important differences: Karmeier et al. (2005) did not investigate the effect of VS coupling, instead focusing on the effects of integration time and input correlations. Furthermore, they used a phenomenological model, in contrast to our biophysically plausible model. More recently, Weber et al. (2012) applied generalized linear models to assess the benefits of coupling between two optic flow-processing, spiking neurons of the lobula plate (H1 and Vi) for conveying information about optic flow parameters.

We found that changes in correlation structure were important in improving encoding accuracy in the case of a subpopulation readout (Figure 7). These findings contrast with typical arguments about the benefit and harm of trial-to-trial correlations (Tkčcik et al., 2010). Correlations between VS neurons carry information about the responses of unobserved neurons. Electrical synapses are both strong and fast in their effect on subthreshold dynamics relative to their chemical counterparts (Xiao et al., 2013). They are therefore well-suited for increasing the coverage of a parameter, reducing variability and introducing correlations.

Many previous theoretical studies examined how changes in neuronal response statistics, such as correlations or tuning curves, impact coding (Barlow, 1961; Sompolinsky et al., 2001; Seriés et al., 2004; Averbeck et al., 2006b; Josić et al., 2009; Ecker et al., 2011; Latham and Roudi, 2011). This can give valuable insights into how coding is affected by aspects of the neural population response. However, correlations and tuning curves are not intrinsic properties of a population response that can be changed arbitrarily (Shea-Brown et al., 2008; Beck et al., 2012). To examine how the statistics of neuronal activity affect coding, it is therefore important to consider realistic networks with realistic inputs.

A key advantage of our approach is that we made no *a priori* assumptions about the VS population response. In particular, we made no assumptions about how the joint activity of VS neurons encodes the axis of rotation. Rather, we considered the responses of a biophysically realistic model to various stimuli. The spatiotemporal structure of the input, and the properties of the VS network fully determined its responses, allowing us to characterize the best estimate of the stimulus available to the animal (Graf et al., 2011).

Our results open a number of avenues for future research: We used temporal averages as a readout of the VS population responses. However, downstream DNOVS neurons are not perfect integrators. DNOVS 2 is a spiking neuron, introducing a strong nonlinearity into the processing pathway. Further, the effect of interactions with other neurons of the lobula plate should be investigated (Borst and Weber, 2011). We also did not attempt to examine the impact of correlations beyond second order. Application of maximum-entropy approaches could help address this topic (Jaynes, 1957).

The aerial performance of flies is unmatched in nature and technology (Frye and Dickinson, 2001). Understanding how a small set of neurons in the fly removes irrelevant variability to extract behaviorally relevant information can provide insight into the implementations in more complex organisms.

Our results in the fly VS network are suggestive of a general principle: Coupling between neurons allows for near optimal readouts from a subpopulation. Correlations between the responses introduced by coupling can lead to ‘information democracy’ where even small subpopulations carry nearly as much information about the stimulus as the entire population. In a neuronal network which encodes a behaviorally relevant parameter, coupling can thus allow each neuron to represent a greater extent of the parameter space. When downstream targets extract information about this parameter from relatively few neuronal projections, correlations between responses can be highly beneficial (Stevenson et al., 2012). While details of our study are particular to the fly visual system, the main ideas are likely to apply across organisms and modalities.

## Acknowledgements

We thank Drs. Alexander Borst and Yishai Elyada for generously providing their computer code for the VS network model, and Elisabeth Hopp for comments on the manuscript. We thank Edgar Gabriel (Department of Computer Science at the University of Houston) and the Texas Learning and Computation Center for access to computer clusters. This work was supported by NSF grants DMS-0817649, DMS-1122094, a Texas ARP/ATP award to KJ, as well as a Humboldt Research Award, NSF grant DMS 1120952 and NIH grant MH065339 to FG.

